# A comparison of methods to suppress electrocardiographic artifacts in local field potential recordings

**DOI:** 10.1101/2022.06.17.496567

**Authors:** M.J. Stam, B.C.M. van Wijk, P. Sharma, M. Beudel, D.A. Piña-Fuentes, R.M.A. de Bie, P.R. Schuurman, W.-J. Neumann, A.W.G. Buijink

## Abstract

**Objective:** Sensing-enabled neurostimulators for deep brain stimulation (DBS) therapy record neural activity directly from the stimulating electrodes in the form of local field potentials (LFPs). However, these LFPs are often contaminated with electrocardiographic (ECG) artifacts that impede the detection of physiomarkers for adaptive DBS research. This study systematically compared the ability of different ECG suppression methods to recover disease-specific electrical brain activity from ECG-contaminated LFPs.

**Approach:** Three ECG suppression methods were evaluated: (1) QRS interpolation of the Perceive toolbox, (2) four variants of a template subtraction method, and (3) sixteen variants of a singular value decomposition (SVD) method. The performance of these methods was examined using LFPs recorded with the Medtronic PerceptTM PC system from the subthalamic nucleus in nine patients with Parkinson’s disease while stimulation was turned off (“OFF-DBS”; anode disconnected) and while stimulation was turned on at 0 mA (“ON-DBS 0 mA”; anode connected). In addition, ECG-contaminated LFPs were simulated by scaling a co-recorded external ECG signal and adding it to the OFF-DBS LFPs.

**Main Results:** ECG artifacts were present in 10 out of 18 ON-DBS 0 mA recordings. All ECG suppression methods were able to drastically reduce the percent difference of beta band (13 – 35 Hz) spectral power and at least partly recover the beta peak and beta burst dynamics. Using predetermined R-peaks improved the performance of the ECG suppression methods. Lengthening the time window around the R-peaks resulted in stronger reduction in artifact-induced beta band power but at an increased risk of flattening the beta peak and loss of beta burst dynamics.

**Significance:** The SVD method formed the preferred trade-off between artifact cleaning and signal loss, as long as its parameter settings (time window around the R-peaks; number of components) are adequately chosen.

## Introduction

Deep brain stimulation (DBS) is a well-established therapeutic option for several neurological and psychiatric disorders, such as Parkinson’s disease (PD), essential tremor, and obsessive compulsive disorder (Krauss et al., 2020; Lozano & Lipsman, 2013). In current DBS therapy, stimulation parameters are adjusted based on an assessment of the patient’s symptom severity. The chosen parameters are then used to deliver stimulation continuously throughout the day, while patients might experience strong fluctuations in experienced motor impairment and therapy-induced side effects. The recent development of sensing-enabled neurostimulators opens up the possibility to record brain activity directly from the stimulating electrode in the form of local field potential (LFP) signals (Swinnen et al., 2022). So-called ‘physiomarkers’ from these LFPs have been successfully used in research studies to automatically adjust stimulation settings, referred to as “adaptive DBS” or “closed-loop DBS” (Beudel et al., 2018; Little et al., 2013; Tinkhauser et al., 2017). In PD, high-amplitude beta (13-35 Hz) oscillations within the subthalamic nucleus (STN) are known to be correlated with the severity of motor symptoms (rigidity and bradykinesia) (Beudel et al., 2017; Neumann et al., 2016; Piña-Fuentes et al., 2019; van Wijk et al., 2016). Crucially, pathological beta oscillations are not continuously increased in amplitude but come in so-called “bursts” of different durations and amplitudes that are reflective of concurrent motor impairment (Lofredi et al., 2022; Tinkhauser et al., 2017). In theory, the efficacy of DBS therapy might be improved by automatically adjusting stimulation to the presence of physiomarkers, such as beta oscillations, in the LFP signal (Habets et al., 2018; Little & Brown, 2020).

The first FDA-approved sensing-enabled neurostimulator, the Medtronic PerceptTM PC, was introduced in 2020 (Jimenez-Shahed, 2021). Previous studies using sensing-enabled neurostimulators showed that it can be difficult to record LFPs of good quality due to electrocardiographic (ECG) artifacts (Anidi et al., 2018; Hell et al., 2018; Neumann et al., 2021; Quinn et al., 2015; Swann et al., 2017). Preliminary studies using the PerceptTM PC showed that ECG artifacts are particularly present in recordings in monopolar stimulation mode (or during calibration test), because the impulse generator (IPG) is both the ground for the recordings and the anode for the stimulation settings (Neumann et al., 2021; Thenaisie et al., 2021). For the PerceptTM PC, monopolar stimulation can be applied by activating the “ON-DBS” status setting. LFPs are concurrently recorded using the “BrainSense Streaming” mode, either without delivering current by setting the stimulation amplitude to 0 mA, or while delivering current by increasing stimulation amplitude. Using the so-called “BrainSense Survey” mode, the monopolar stimulation is turned off (subsequently referred to “OFF-DBS”), the IPG is uncoupled from the stimulation contact and LFPs can be recorded for up to approximately 20 seconds (Jimenez-Shahed, 2021). Using this recording mode, ECG contamination is very rare (Neumann et al., 2021; Thenaisie et al., 2021), which therefore serves as a control condition in our study.

As previously reported, a significant number of LFPs recorded with implantable devices are contaminated by ECG artifacts. Without further processing, these signals are not usable for research or clinical purposes. To adequately deal with this issue, robust ECG artifact suppression methods must be developed. Previous studies have adopted various methods to attenuate ECG artifacts in LFP signals (Canessa et al., 2020; Chen et al., 2021; Thenaisie et al., 2021). The performance of different approaches have recently been compared, which all were found to successfully remove ECG artifacts (Hammer et al., 2022). Nevertheless, there remains an unmet need in the field of clinical brain computer interfaces for DBS (a) to corroborate the results with a larger sample size, (b) to objectively assess the approaches by means of simulations, and (c) to evaluate the actual impact of the methods on the physiomarkers themselves (e.g., beta power and beta bursts distribution). To this end, we compared several ECG suppression methods at the stage of signal processing and assessed their ability to preserve important physiomarkers of PD. The suppression methods were tested on LFPs with real ECG artifacts and validated by applying them to LFPs with simulated ECG artifacts.

## Materials and methods

### Participants

We included LFP recordings from nine patients with Parkinson’s disease (Supplementary Materials; Table S1). All patients were bilaterally implanted in the subthalamic nucleus (STN) between 2013-2016 (electrode lead model 3389, Medtronic, Minneapolis, MN). Due to battery depletion, the pulse generator was replaced with the PerceptTM PC neurostimulator (Medtronic, Minneapolis, USA) in the context of standard clinical care between September 2020 and March 2021. The recordings were performed within six months after replacement surgery. This study was approved by the local ethics committee and was carried out in accordance with the Declaration of Helsinki. Informed written consent was received from all patients.

### Data Acquisition

During the experiment, patients lay on a bed with the headrest in 45 degrees, or sat comfortably in a chair with the arms supported, and were instructed not to talk or sleep during recordings. At the start of the experiment, stimulation was turned off for at least 15 minutes. Subsequently, the so-called “BrainSense Survey” mode was used to record LFP signals of approximately 20 seconds (5288 samples) with a sampling frequency of 250 Hz from all contact pairs (“OFF-DBS”, bipolar recordings for both sides using contact combinations 0-2, 1-3 and 0-3). After these twenty-second “OFF-DBS” LFP recordings, the stimulation was turned on with a stimulation amplitude of 0 mA in the “BrainSense Streaming” mode (“ON-DBS 0 mA”). LFP signals were recorded at a sampling frequency of 250 Hz. Contact points surrounding the one used for clinical stimulation were used for bipolar recordings from both “BrainSense Survey” and “BrainSense Streaming” data. LFP signals were recorded for approximately 60 seconds while the patients were at rest. The ON-DBS 0 mA LFP signals were visually inspected for the presence of ECG artifacts (by B.W. and M.S.), and the R-peaks of the ECG artifact were visually identified (by P.S. and M.S.). For accurate comparisons between the ON-DBS 0 mA LFP signals and the OFF-DBS LFP signals, a similar recording length of 20 seconds of both signals was used for further analysis. By default, the PerceptTM PC applies a 100 Hz low-pass filter and a 1 Hz high-pass filter during recordings. Raw LFPs were amplified by 250x, buffered on the device, and streamed wirelessly to the PerceptTM PC tablet programmer. This data was exported as JSON-file and processed offline using custom-made scripts, the open-source Perceive toolbox (www.github.com/neuromodulation/perceive/), and the FieldTrip Toolbox for EEG/MEG-analysis (http://fieldtriptoolbox.org, (Oostenveld et al., 2011)) in MATLAB (version R2020b, The MathWorks, Inc., Natick, MA, USA). A simultaneous bipolar ECG signal was recorded using two electrodes placed on the right and left shoulder (locations: RA and LA of ECG 12 Lead Placement; TMSi Porti amplifier: monopolar, average reference, anti-aliasing low-pass filter with a cut-off frequency of 500 Hz and sampling frequency of 2048 Hz, TMSi, The Netherlands). A band-stop (notch) filter was applied to remove the power line interference (between 49-51 Hz) and its harmonics (99-101 Hz). Synchronization of the LFP and ECG signals was done by slightly ramping up and down the stimulation amplitude at the start of each recording. The induced stimulation artifacts visible in both signals (i.e., LFP and external ECG) were aligned at the stage of post-processing in order to synchronize LFP and ECG recordings. The ECG signals were subsequently down-sampled to 250 Hz to match the sampling rate of the LFP signals. This method has been used previously (Steiner et al., 2017; Thenaisie et al., 2021).

### Artifact Simulation

Recording in “BrainSense Survey” mode resulted in OFF-DBS LFP signals that were free from ECG artifacts. The first 20 seconds (5288 samples at 250 Hz) of each externally measured ECG signal were used to create time series with varying degree of ECG contamination (Abbaspour & Fallah, 2014). The recorded external ECG signals were scaled to meet eleven levels of simulated ECG artifact and were added to the OFF-DBS LFP signals. For the lowest level of ECG artifact simulation, the R-peaks were at the level of the mean amplitude envelope of the OFF-DBS LFP as determined via the Hilbert transform (0%). The highest level of ECG artifact simulation was set such that the average percent difference of beta band power between the simulated LFP signal and the original OFF-DBS LFP signal was similar to the percent difference of beta band power as found for the ON-DBS 0 mA with ECG artifacts. This implied that the highest level of ECG artifact was simulated by multiplying the lowest ECG artifact level by eleven (100%). Levels 10 to 90% of ECG artifact were obtained by multiplying the lowest ECG artifact level by two to ten, respectively. An example of the simulated ECG artifacts is shown in Figure S1 in the Supplementary Materials.

### ECG Suppression Methods

Three methods of ECG artifact suppression were assessed: QRS interpolation as implemented in the Perceive toolbox (Perceive, https://github.com/neuromodulation/perceive (Neumann et al., 2021)), a custom template subtraction method (Template), and a singular value decomposition (SVD) method.

#### I. Perceive Toolbox Method

*Perceive* is a versatile low-weight MATLAB based toolbox for json file conversion, data standardization and preprocessing of PerceptTM PC data. One of the functions is ECG cleaning through QRS interpolation that is available through the online repository of the Perceive toolbox (Neumann et al., 2021). The QRS complex is the most prominent peak in the ECG artifact and corresponds to the ventricular depolarization phase. This method identifies R-peaks through a series of operations:

1. segmentation of the LFP signal into arbitrary 1s epochs;
2. alignment of the epochs using cross-correlation followed by averaging to obtain a first template;
3. search for the highest and lowest peaks to crop the template to the putative ECG artifact period;
4. perform temporal correlation between the first ECG template and the original signal with a sliding window of 1 sample;
5. perform adaptive thresholding of the correlation coefficients to maximize the number of identified peaks with an interpeak interval that is typical for ECG (55-120 bpm);
6. create a second ECG template by averaging over all epochs created around the detected ECG peaks (50 ms before- and 100 ms after the peak location);
7. repeat steps 3, 4, and 5 to obtain a second threshold for the correlation coefficient; The correlation coefficients that cross the second threshold are the locations of the final QRS artifact.

The peak detection algorithm of the Perceive toolbox identifies LFP signals as being contaminated by ECG artifact if the heart rate was between 50 and 120 bpm, if, on average, an R-peak was found once every two seconds, if the difference between the lowest and highest peak found in step 3 was at least 0.075 mV, and if the final ECG peaks were at least 20% higher than the inter-peak interval. The artifact is subsequently removed by interpolation, where the QRS complex (156 ms) is replaced by mirrored-padding the signal immediately preceding and following the identified QRS complex using a window of the final ECG template length surrounding the peak location.

#### II. Template Subtraction Method

The template subtraction method subtracts the average ECG-waveform across all R-peaks from the continuous time series. It can be broken down into four steps (see Figure 1). The first step is to identify the locations of the QRS complexes of the ECG artifact. Two different algorithms of R-peak detection were developed and tested:

**Figure 1.**
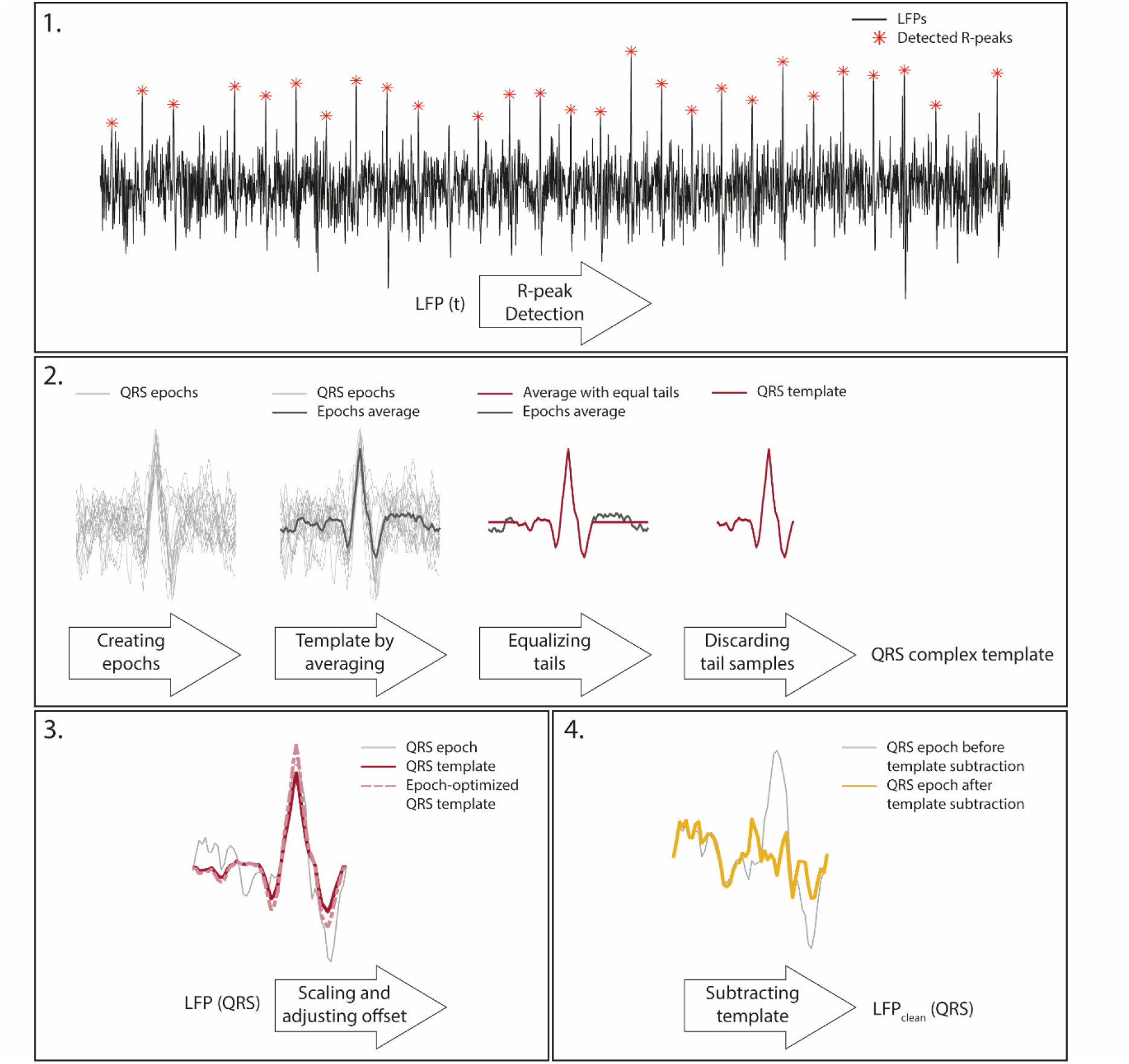
Schematic overview of the template subtraction method. In this example, the QRS time window is visualized. Step 1: the R-peaks (red asterisks) of the ECG artifact in the recorded LFP signal (LFP(t); thin black line) are detected; Step 2: epochs around R-peaks are created (LFP(QRS); thin grey lines) and averaged to create the initial template (thick black line). The tails of the initial template (maximum 30 samples) were searched for two samples whose values had the smallest difference. These samples were set as the start and end points of the template (thick red line); Step 3: per epoch (LFP(QRS); thin grey line), the final template (thick red line) is scaled and the offset between the original R-peak and the template is corrected (pink dashed line); Step 4: the optimized template is subtracted from the original epoch (LFP(QRS); thin grey line) to obtain a cleaned LFP signal (LFPclean(QRS); thick yellow line).

1a. *Peak detection using the LFP signal (Template (LFP))*: the LFP signal was z-scored (X-μ/σ) over the entire recording. Subsequently, the build-in MATLAB function *findpeaks* was used to search for R-peaks with a specific height (at least two-and-a-half times the standard deviation of the entire time series) and at a specific inter-peak distance (minimally 500 ms). The algorithm accounts for a negative QRS complex by repeating this procedure after multiplying the signal with -1. For both orientations of the LFP signal, the values of the peaks were averaged and the peaks with the highest mean determined the orientation of the QRS complexes, provided that at least the same amount of peaks were detected.

1b. *Peak detection using the ECG signal (Template (ECG))*: the externally recorded ECG signal was used to predetermine the timestamps of the R-peaks. The ECG signal was z-scored (X-μ/σ) over the entire recording and the same algorithm as described in *1a* was used to search for R-peaks. Subsequently, the LFP signal was searched for peaks using MATLAB functions *min* and *max* in a window of 20 samples prior and after the R-peaks found in the ECG signal. This window was adopted in order to account for inaccuracies in the synchronization of the LFP- and ECG signals. The absolute values of the peaks found with *min* and *max* were averaged and the peaks with the highest absolute mean determined the orientation of the QRS complexes.

To avoid LFP signals without visual ECG artifacts to inaccurately be cleaned, the template peak detection algorithm only identified LFP signals as contaminated with ECG artifacts if at least one R-peak every three seconds and a heart rate of at least 40 beats per minute was detected.

The second step is to generate a template. Epochs were created by selecting a time window around each detected R-peak. Two different time windows were used to create either QRS- or PQRST-complex templates to assess the effect of epoch length on artifact-suppression performance.

2a. *QRS complex template generation*: QRS complex epochs were created by taking 200 ms (50 samples) before and after each detected R-peak in the LFP signal. Epochs were averaged over all R-peaks to create the initial template. Subsequently, the Q- and S-peaks were detected with the MATLAB function *findpeaks.* Both outer sides of the Q- and S-peaks (tails with a maximum of 30 samples) were searched for two samples whose values had the smallest difference. These samples were set as the start and end points of the template. In case of dissimilar values, the offset was corrected by changing the point with the lower value to the value of the higher point. Equal values for start and end points avoid any introduced offsets between the corrected epochs and neighbouring data points.

2b. *PQRST complex template generation:* PQRST complex epochs were created by taking 250 ms before and 400 ms after the R-peaks. The size of the window ensured that the P- and T-waves were included, thereby accepting the risk of overlapping epochs in case of a high heart rate. Again, the epochs were averaged over all R-peaks to create the template. The first and last 5 ms were searched for equal values to determine the start and end values, in the same way as described above.

3. *Template optimization*: for each epoch in the original LFP signal the final template (QRS or PQRST; panel A in Figure S2 and S3 of Supplementary Materials, respectively) was optimized by adjusting the scale and offset parameters such that the sum of squared error between the original LFP signal and optimized template is minimized, using the build-in MATLAB function *lsqnonlin*:

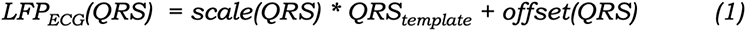

4. *Template subtraction:* this epoch-specific optimized template, LFPECG(QRS/PQRST), was then subtracted from the corresponding epoch in the original LFP signal, LFP(QRS/PQRST), to obtain the LFP signal at the epochs without ECG artifact: LFPclean(QRS/PQRST):

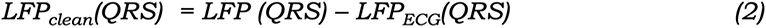

#### III. Singular Value Decomposition Method

Singular value decomposition (SVD) was applied to accommodate variations in ECG waveform shape across epochs. It was implemented similarly to previous studies that used this method for removal of ECG- and spike-related artifacts from DBS-LFP recordings (Canessa et al., 2020; Hammer et al., 2022; Scheller et al., 2019; Thenaisie et al., 2021). The same peak detection algorithms (SVD (LFP) and SVD (ECG)) and time windows (SVDQRS and SVDPQRST) were used as described for the template subtraction method to create epochs. The matrix of *M* time points x *N* epochs was subsequently decomposed into a set of singular vectors and corresponding singular values. The ECG artifact for each epoch was reconstructed from the components with largest singular values and was subtracted from the original time series. Again, to avoid any introduced offsets between the corrected epochs and neighbouring data points, per epoch, the first and last 4 ms of the reconstructed ECG artifact were searched for two samples whose values had the smallest difference, and were set equal in case of dissimilar values (in the same way as described for the template subtraction method). Per epoch, the offset parameter was adjusted such that the sum of squared error between the original LFP signal and reconstructed ECG artifact was minimized (application of Formula 1, only for offset). In general, the outcome of this method is expected to depend on the number of selected components. A larger number of components can capture the ECG artifact more completely but comes with the risk of removing LFP signal features of interest. We investigated how strongly this trade-off influenced the performance of the SVD method by comparing the use of one (SVD1), two (SVD2), three (SVD3), or four (SVD4) components. The cumulative percentage energy of the components is shown in Table S2 in the Supplementary Materials. As an example, the SVD1-4PQRST (LFP) components found for the left STN-LFP signal of Sub04 is shown in Figure S4 in the Supplementary Materials. A visualisation of the resulting ECG artifact of the SVD1-4 (LFP) that was subtracted at the epochs of the original time series is shown in panels B-E of Figures S1 (for QRS time window) and S2 (for PQRST time window) in Supplementary Materials.

### Offline Spectral Analysis

Power spectral densities (PSDs) of LFP signals were calculated both before and after applying the ECG suppression methods using the Welch method (MATLAB function *pwelch*) with 1s Hamming windows and 50% overlap. The relative PSD was obtained by normalizing each power spectrum to the total summed power over the 1-49 and 51- 99 Hz frequency ranges of all PSDs (OFF-DBS LFPs and ON-DBS 0 mA before and after applying the ECG suppression methods). The 0-1 and 49-51 Hz ranges are omitted in order to avoid contamination by movement artifact and mains noise, respectively. For the ON-DBS 0 mA LFPs without ECG artifact, the PSD was only normalized over the summed power of the OFF-DBS LFPs and ON-DBS 0 mA before ECG suppression. In addition to the relative PSDs, the average power within the beta band (13 – 35 Hz) was calculated for each original and cleaned signal. These values were subsequently expressed as a percent difference of power between ON-DBS 0 mA LFPs without ECG artifact and corresponding OFF-DBS LFPs, between cleaned ON-DBS 0 mA LFPs and corresponding OFF-DBS LFPs, and between cleaned simulated signals and their original OFF-DBS LFPs. Grand-average differences and standard deviation were computed across hemispheres and patients.

### Beta Bursts Occurrence

For the beta burst analysis, only OFF-DBS LFP signals containing a clear beta peak were selected. The original and simulated LFP signals were decomposed using Wavelet transformation (*ft_specest_wavelet* function in Fieldtrip - Morlet Wavelet, width = 7, gwidth = 3; Oostenveld et al., 2011) into frequency components between 13 and 35 Hz with a frequency resolution of 1 Hz. For each LFP signal, a threshold was set at the 75th percentile of the averaged beta band power. Each data point at which the mean time-evolved beta-amplitude exceeded the 75th percentile threshold was selected as (part of) a beta burst (Tinkhauser et al., 2017). The minimum duration of a beta burst was set at 150 ms. Each sample of the simulated signal was then assigned a label after comparison with the same sample in the original OFF-DBS LFP signal: true burst (true positive, TP), true non-burst (true negative, TN), false burst (false positive, FP), or false non-burst (false negative, FN). The false alarm rate and hit rate were calculated (Formula 3), after which the performance of the ECG suppression methods was compared using sensitivity index *d’* and bias criterion *c* (Stanislaw & Todorov, 1999) using the MATLAB function *dprime* (Böckmann-Barthel, n.d.). A value of *d’* = 3 is close to perfect performance; a value of *d’* = 0 is chance (“guessing”) performance. In signal detection theory, a positive *c* value reflects a bias towards high hit rates and false alarm rates (Albrecht & Mattler, 2012).

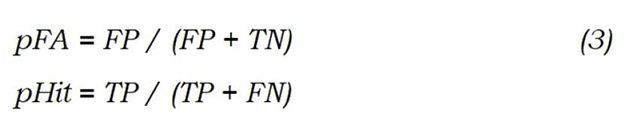

### Statistical Analyses

The performance of the peak detection algorithms was determined by comparing the number of correctly identified peaks with the actual number of peaks in the externally recorded ECG signal as ground truth. Non-parametric Wilcoxon signed-rank tests were used for these comparisons, as data points were not normally distributed.

Spectral differences were statistically assessed using the percent difference in beta power as outcome measure and a significance threshold of 0.05. The difference between ON-DBS 0 mA LFP signals without visual ECG and OFF-DBS LFP signals were tested using a two-sided one-sample t-test. A Mann-Whitney U-test was used to test for differences between ON-DBS 0 mA LFP signals without ECG artifact and the ON-DBS 0 mA LFP signals with ECG artifact, as assumptions for a parametric test were not met for the latter group. ECG suppression method (Template, SVD1-SVD4), R-peak detection method (using the LFP signal only, or with use of the externally recorded ECG signal), and the length of the time window around the R-peaks (QRS or PQRST) relative to the ON-DBS 0 mA LFP signals were compared using a 5 x 2 x 2 repeated measures ANOVA. In a separate repeated measures one-way ANOVA, the Perceive method was compared to the Template and SVD methods that use the LFP signal to detect R-peaks and the QRS time window. Both ANOVA tests were followed by post-hoc paired two-sample t-tests with Bonferroni adjustment. Friedman ANOVA tests and qualitative comparisons were performed for the simulated data as percent difference values were found to be not normally distributed.

## Results

Nine PD patients were included (Supplementary Materials; Table S1). This resulted in 54 artifact-free OFF-DBS LFP signals (six bipolar recordings per patient based on three contact pairs per electrode) and 18 ON-DBS 0 mA LFP signals (two bipolar recordings per patient based on one contact pair per electrode).

### ECG Artifact Detection

Visual inspection of the ON-DBS 0 mA LFP signals found that ten signals contained an ECG artifact (Kappa’s inter-rater reliability (two raters) = 1). The Perceive algorithm of the QRS interpolation correctly identified six, and the algorithm of the template subtraction method that only uses the LFP signal (Template (LFP)) seven. An example of an ON-DBS 0 mA LFP signal with ECG artifacts is shown in red in Figure 2.

**Figure 2.**
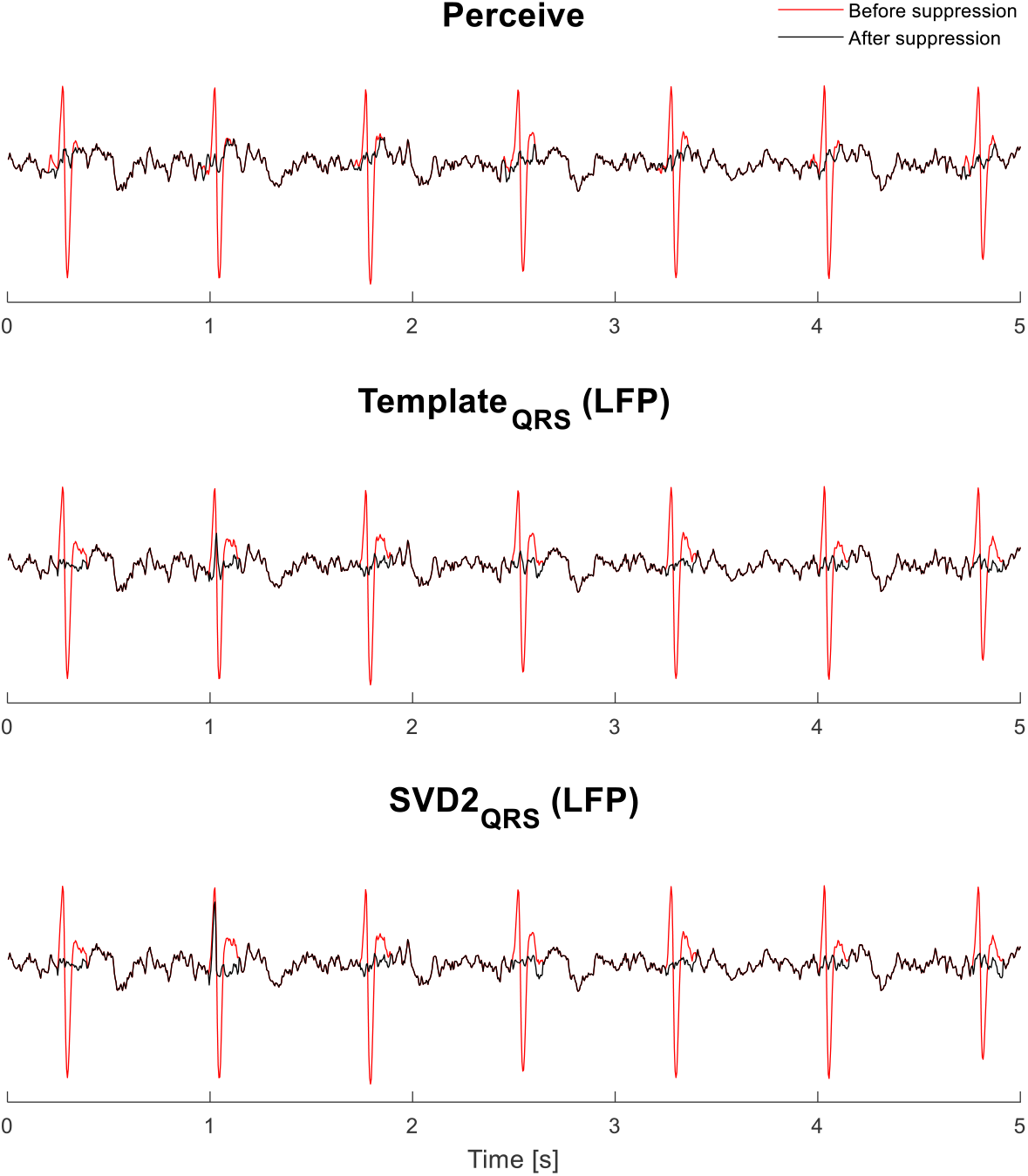
An example of an ON-DBS 0 mA LFP signal with ECG artifact. Five seconds of an ON-DBS 0 mA LFP signal before (red line) and after (black line) applying the ECG suppression methods.

In five ON-DBS 0 mA LFPs, the R-peaks of the ECG artifact lay close to the level of neural activity. For these signals, no consensus could be reached between researchers (P.S. and M.S.) when visually identifying the R-peaks of the ECG artifact. Therefore, the externally recorded ECG signal was used to confirm the timestamps in the LFPs at which the R-peaks occurred. The number of identified R-peaks in the ON-DBS 0 mA LFP signals (60 seconds) using the peak detection algorithm of the Perceive toolbox method (mean = 68.5, sd = 19.5) as well as the number of R-peaks identified by the algorithm of the Template (LFP) method (mean = 71.1, sd = 16.5) significantly differed from the number of R-peaks found while using the externally recorded ECG signal (mean = 79.0, sd = 13.7; Perceive toolbox method *Z* = -2.366, *p* = .018 and Template method *Z* = -2.023, *p* = .043). The performance of the algorithms of the Perceive toolbox method and Template (LFP) method was similar (*Z* = -1.261, *p* = .207).

### Spectral Analysis

Accurate comparisons between the ON-DBS 0 mA LFP signals and the OFF-DBS LFP signals was assured by using a similar recording length of 20 seconds for both signals. The averaged relative PSD of the ON-DBS 0 mA LFP signals without ECG artifacts and the corresponding OFF-DBS LFPs showed a clear beta peak (Figure 3A). The beta peak was concealed in ON-DBS 0 mA LFPs contaminated with ECG artifacts (Figure 3B).

**Figure 3.**
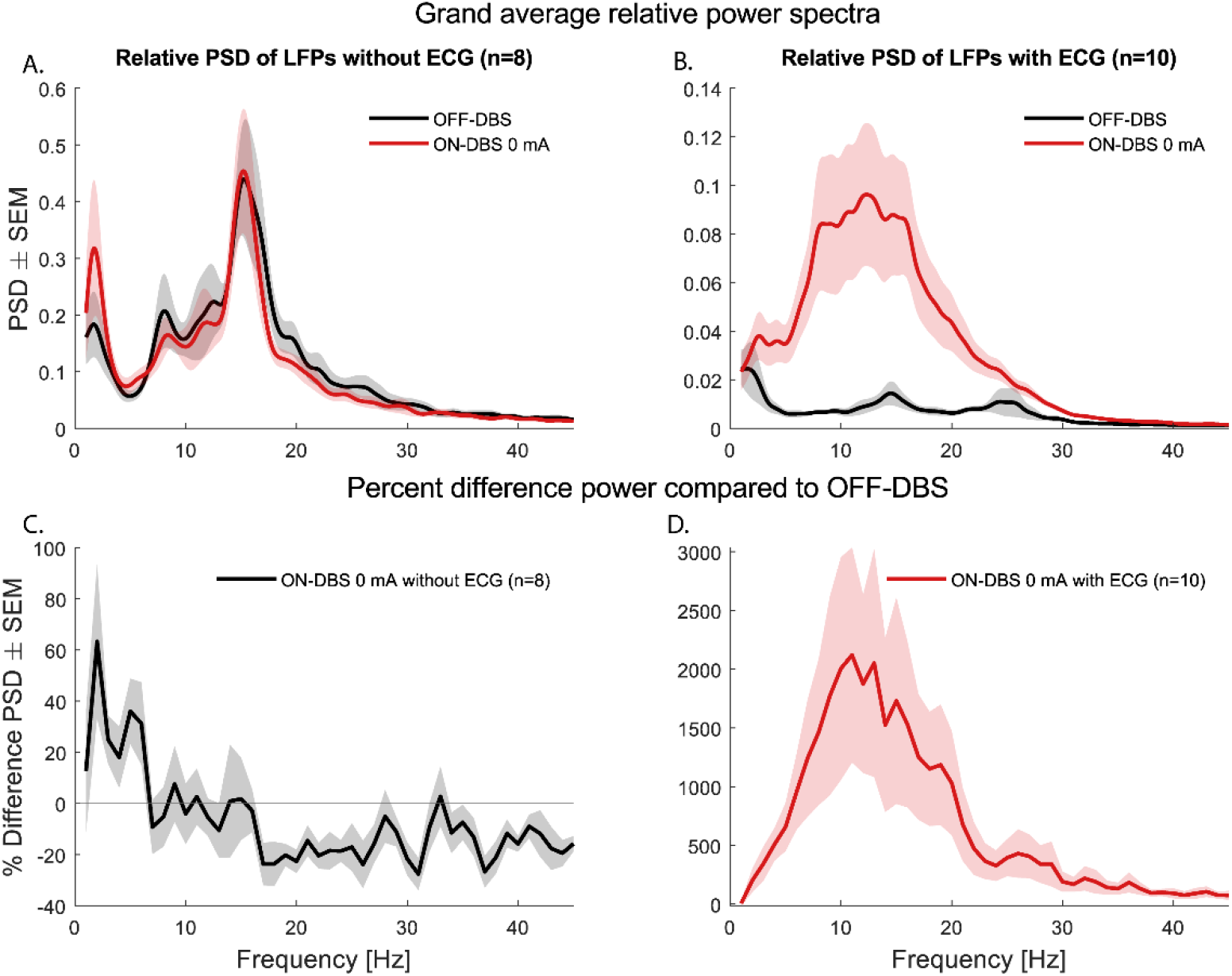
The effect of ECG artifact in the frequency domain before applying the ECG suppression methods. Panels A-B: the grand average relative PSD of ON-DBS 0 mA LFPs (red line) without ECG artifact and corresponding OFF-DBS LFPs (black line) (A) and the grand average relative PSD of ON-DBS 0 mA LFPs with ECG artifact and corresponding OFF-DBS LFPs (B). Panels C-D: average percent spectral power difference for the ON-DBS 0 mA LFP signals without (black line; C) and with (red line; D) ECG artifacts relative to the corresponding OFF-DBS LFP signals.

The average percent difference of spectral power in the beta band (13 – 35 Hz) of ON-DBS 0 mA LFP signals without visual ECG artifacts (n = 8) significantly differed (mean difference = -14.2%, sd = 10.8%) from the OFF-DBS LFP signals recorded from the same electrode pairs (*t*(7) = -3.716, *p* = .007) (Figure 3C; Table 1). In LFPs contaminated with ECG artifacts (n = 10) the difference in beta power drastically increased (mean difference = 689.2%, sd = 878.0%) leading to both qualitative and quantitative spectral changes (*U* = 0, *p* < .001) (Figure 3D; Table 1).

**Table 1.**
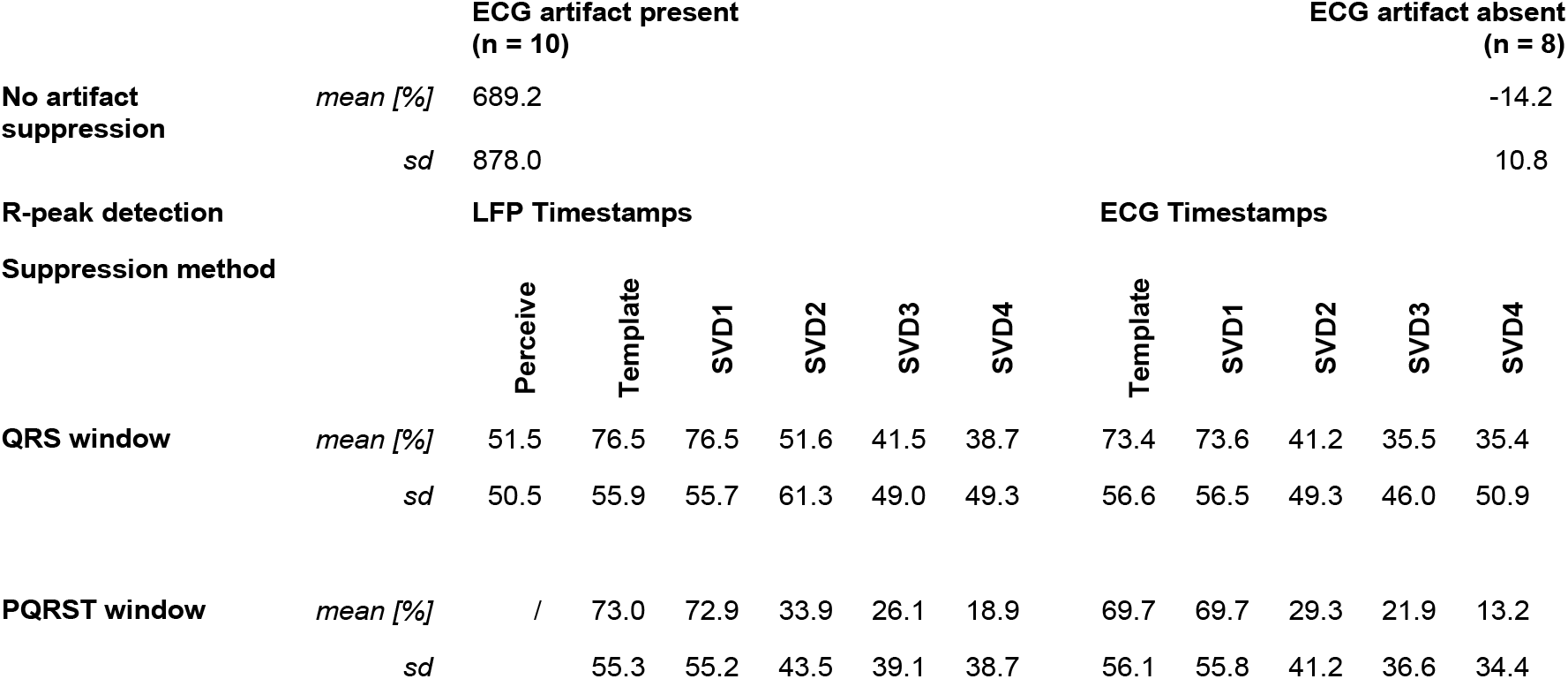
Average percent difference of beta band power between the ON-DBS 0 mA LFP signal and corresponding OFF-DBS LFP signal, before and after applying the ECG suppression methods.

All ECG suppression methods were able to recover the beta peak in the ON-DBS 0 mA LFPs contaminated with ECG artifacts. However, Figure 4C-D shows that the beta peak is least preserved when applying the SVD3 or SVD4 method with the PQRST time window.

**Figure 4.**
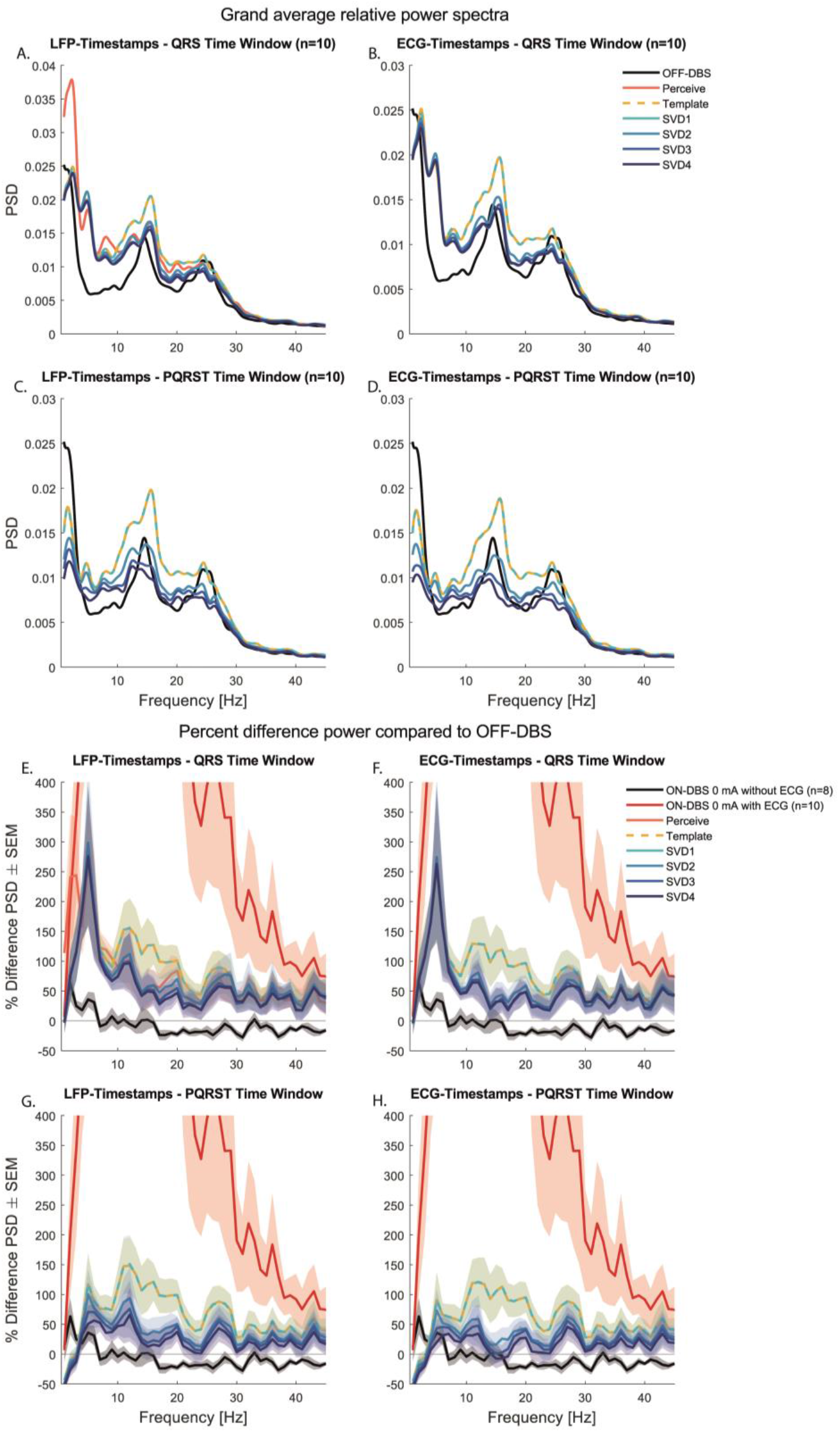
The effect the ECG artifact suppression methods in frequency domain. Panels A-D: the grand average relative PSD of ON-DBS 0 mA LFPs with ECG artifact after applying the ECG suppression methods and corresponding OFF-DBS LFPs (black line). Panels E-H: Average percent spectral power difference for the ON-DBS 0 mA LFP signals with (red line) and without (black line) ECG artifact relative to the corresponding OFF-DBS LFP signals. Each color represents the difference in spectral power after applying the indicated ECG suppression method to the ON-DBS 0 mA LFP signals with ECG artifact.

The average spectral differences for the ON-DBS 0 mA with ECG artifacts before and after applying the ECG suppression methods are visualized in Figure 4E-H. In these panels, the performance of the suppression methods can be compared with the spectral differences for the ON-DBS 0 mA without ECG artifacts, similarly to the Figure 3C. The percent difference for the averaged beta band power after applying the ECG suppression methods is listed in Table 1. These results are also graphically visualized in Figure 5.

**Figure 5.**
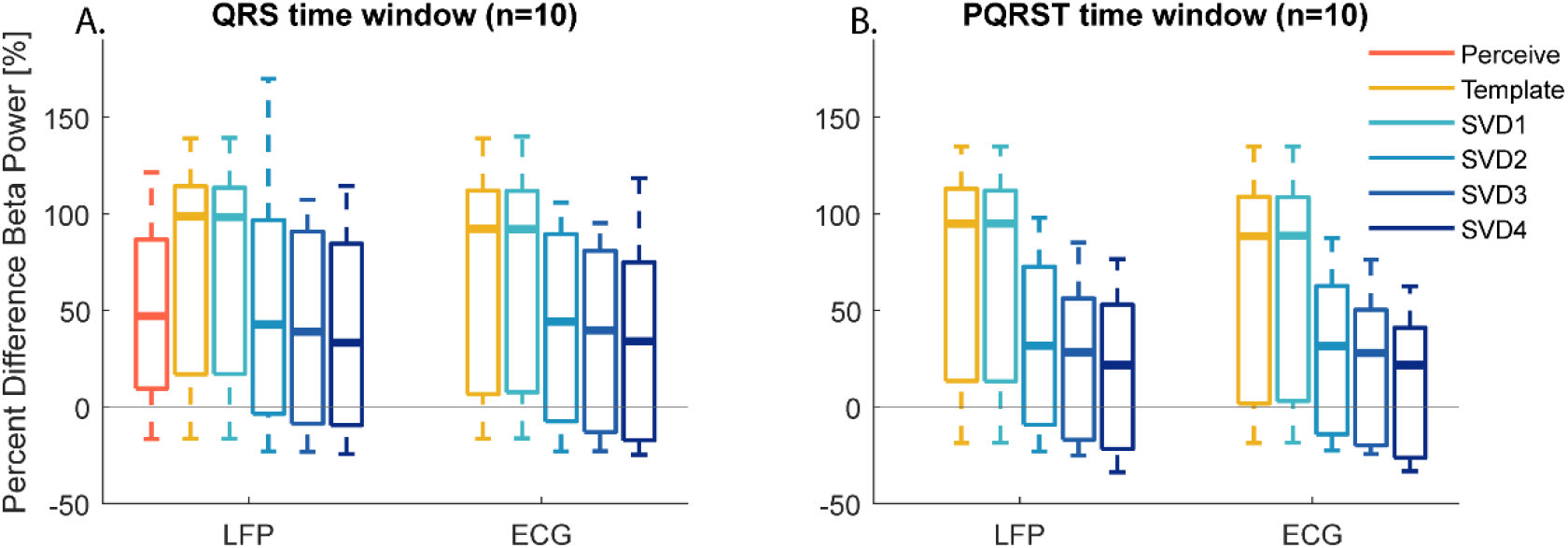
Average percent beta band power difference after applying the ECG suppression methods. Average percent difference of beta band power for the ON-DBS 0 mA LFP signals with ECG artifact (relative to the OFF-DBS LFP signals of the same electrode pair) after applying all ECG suppression methods. The whiskers indicate the lower and upper adjacent values (i.e. the smallest or largest, respectively, data point that is not an outlier in the plot; Formula S1 in Supplementary Materials).

A significant effect of method was found (*F*(5,5) = 14.147, *p* = .006) when comparing the average percent difference of beta power of the six ECG suppression methods (Perceive, and Template and SVD methods that only use the LFP signal to detect R-peaks and the QRS time window) (left cluster of boxplots in Figure 5A). The 5 x 2 x 2 repeated measures ANOVA also showed a significant main effect of method on the average percent difference of beta power after applying the different Template and SVD ECG suppression methods (*F*(4, 6) = 20.953, *p* = .001). Post hoc analysis with a Bonferroni adjustment revealed significant differences between the SVD2 and SVD3 methods (*p* = .026) and between SVD2 and SVD4 methods (*p* < .001). Additionally, artifact-induced beta band power significantly reduced with increasing time window (*F*(1, 9) = 6.277, *p* = .034), and by using the externally recorded ECG signal to detect R-peaks (*F*(1, 9) = 9.805, *p* = .012). There was a statistically significant interaction between the effects of method and time window on average percent difference of beta power (*F*(4, 6) = 4.942, *p* = .042). Figure 5 shows that the longer the time window, the larger the differences between the methods. Other interaction effects were not significant.

### Artifact Simulation

Scaling the nine externally recorded ECG signals into eleven levels of simulated ECG artifact and adding them to all 54 OFF-DBS LFP signals, resulted in 5346 simulations of ECG contaminated LFP signals (9 x 11 x 54). The grand-averaged relative PSD of the simulated LFP signals with 0% ECG contamination and the corresponding OFF-DBS LFPs showed a clear beta peak (Figure 6A). The beta peak was concealed in LFPs contaminated with 100% ECG artifacts (Figure 6B).

**Figure 6.**
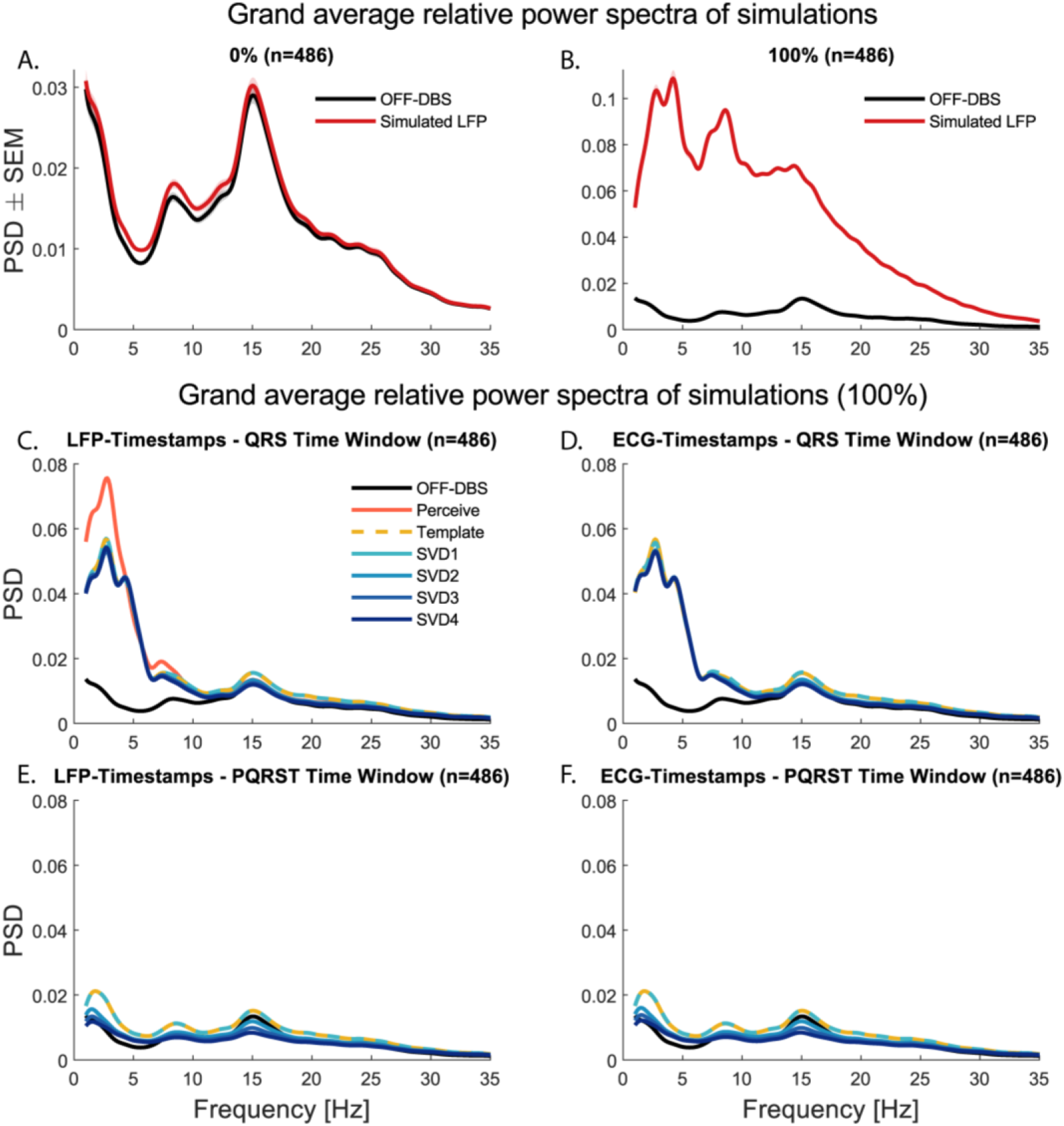
Averaged relative PSDs of the simulated LFPs before and after applying the ECG suppression methods. The grand average relative PSD of simulated LFPs (red line) with 0% ECG contamination and corresponding OFF-DBS LFPs (black line) (A) and the grand average relative PSD of simulated LFPs with 100% ECG artifact (red line) and corresponding OFF-DBS LFPs (black line) (B). Panels C-F: the grand average relative PSD of OFF-DBS LFPs (black line). Each color represents the relative PSD after applying the indicated ECG suppression method.

The ECG suppression methods were applied to the simulated signals to objectively assess their performance on beta band preservation. The relative PSD of the simulated LFPs with 100% ECG contamination after applying the ECG suppression methods confirm that all ECG suppression methods were able to recover the beta peak (Figure 6C-F). Figure 6E-F again show that the beta peak is least preserved when applying the SVD3 or SVD4 method with the PQRST time window.

The average percent difference between the simulated and the original OFF-DBS LFPs for the beta band (13-35 Hz) after applying the ECG suppression methods are shown Table 2 and Figure 7. For these results, all levels of ECG contamination were averaged. Results from the simulations confirmed the pattern of significant (main) effects of method, time window, and peak detection method (Friedman ANOVA all p ≤ .001) as obtained for the ON-DBS 0 mA LFP signals. The SVD4 method with PQRST time window reduced the percent difference of the beta band power most. When dividing the simulations into level of ECG contamination (Supplementary Materials; Figure S5), it becomes apparent that using a longer time window and more components for the SVD method leads to a higher reduction of the percent difference of the beta band power of the simulated signals with high levels of ECG contamination. For simulations with minor ECG artifact, these parameter settings result in negative percent differences.

**Figure 7.**
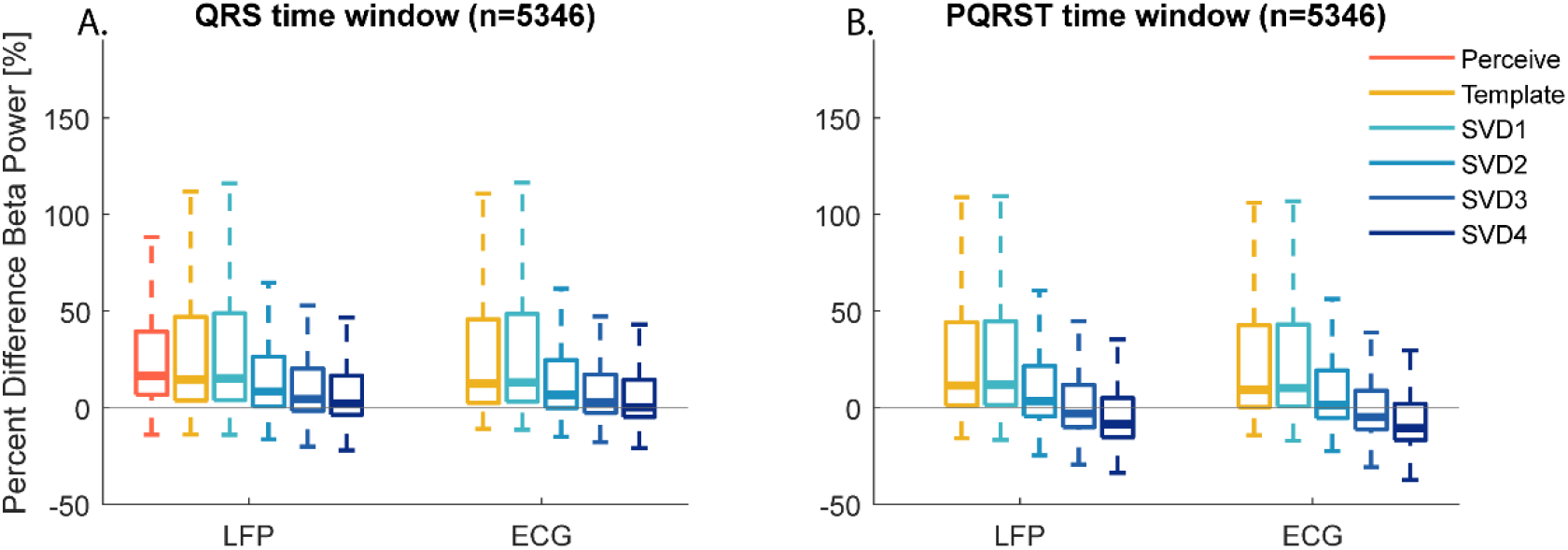
Average percent beta band power difference after applying the ECG suppression methods. Average percent difference of beta band power for all simulation signals relative to the OFF- DBS LFP signals of the same electrode pair after applying all ECG suppression methods. The whiskers indicate the lower and upper adjacent values.

**Table 2.** Percentage difference of beta band power between the simulated LFP signals (n = 5346) and corresponding OFF-DBS LFP signal, before and after applying the ECG suppression methods.

|  |  |  |  |  |  |  |  |  |  |  |  |  |
| --- | --- | --- | --- | --- | --- | --- | --- | --- | --- | --- | --- | --- |
| **No artifact suppression** | *mean [%]* | 287.3 |  |  |  |  |  |  |  |  |  |  |
|  | *sd* | 366.6 |  |  |  |  |  |  |  |  |  |  |
| **R-Peak detection** |  | **LFP Timestamps** |  |  |  |  |  | **ECG Timestamps** |  |  |  |  |
| **Suppression method** |  | **Perceive** | **Template** | **SVD1** | **SVD2** | **SVD3** | **SVD4** | **Template** | **SVD1** | **SVD2** | **SVD3** | **SVD4** |
| **QRS window** | *mean [%]* | 35.9 | 43.6 | 44.8 | 25.8 | 22.5 | 18.5 | 42.4 | 45.5 | 25.8 | 20.9 | 18.1 |
|  | *sd* | 62.7 | 79.5 | 80.6 | 56.7 | 58.4 | 53.0 | 79.7 | 87.1 | 61.4 | 58.6 | 57.6 |
| **PQRST window** | *mean [%]* | / | 40.2 | 40.6 | 19.5 | 11.1 | 4.3 | 39.1 | 39.6 | 18.1 | 9.4 | 2.4 |
|  | *sd* |  | 77.4 | 77.6 | 52.9 | 48.2 | 45.0 | 77.5 | 77.6 | 53.2 | 48.2 | 45.3 |

### Beta Bursts Analysis

The PSDs of the 54 OFF-DBS LFP signals were visually inspected for the presence of a beta peak. Seventeen signals were selected, which means 1683 (17 LFP signals * 9 ECG simulations * 11 levels of artifact) simulations were used for the beta burst analysis. On average, the original OFF-DBS LFP signals had 12.1 ± 2.0 bursts (mean ± sd) of 268.2 ± 27.8 (mean ± sd) ms. The number and duration of the bursts found before and after applying the ECG suppression methods are listed in Table S3 in Supplementary Materials. For these results, all levels of ECG contamination were averaged. The distribution of the median beta bursts durations in the original signals and the ECG artifact simulations, both prior and after applying the ECG suppression methods is visualized in Figure 8. Beta burst distributions after applying the ECG suppression methods were qualitatively similar compared to those obtained for the OFF-DBS LFP signals. A slight shift towards relative more bursts at very short and long duration, and fever bursts with intermediate duration seemed to remain.

**Figure 8.**
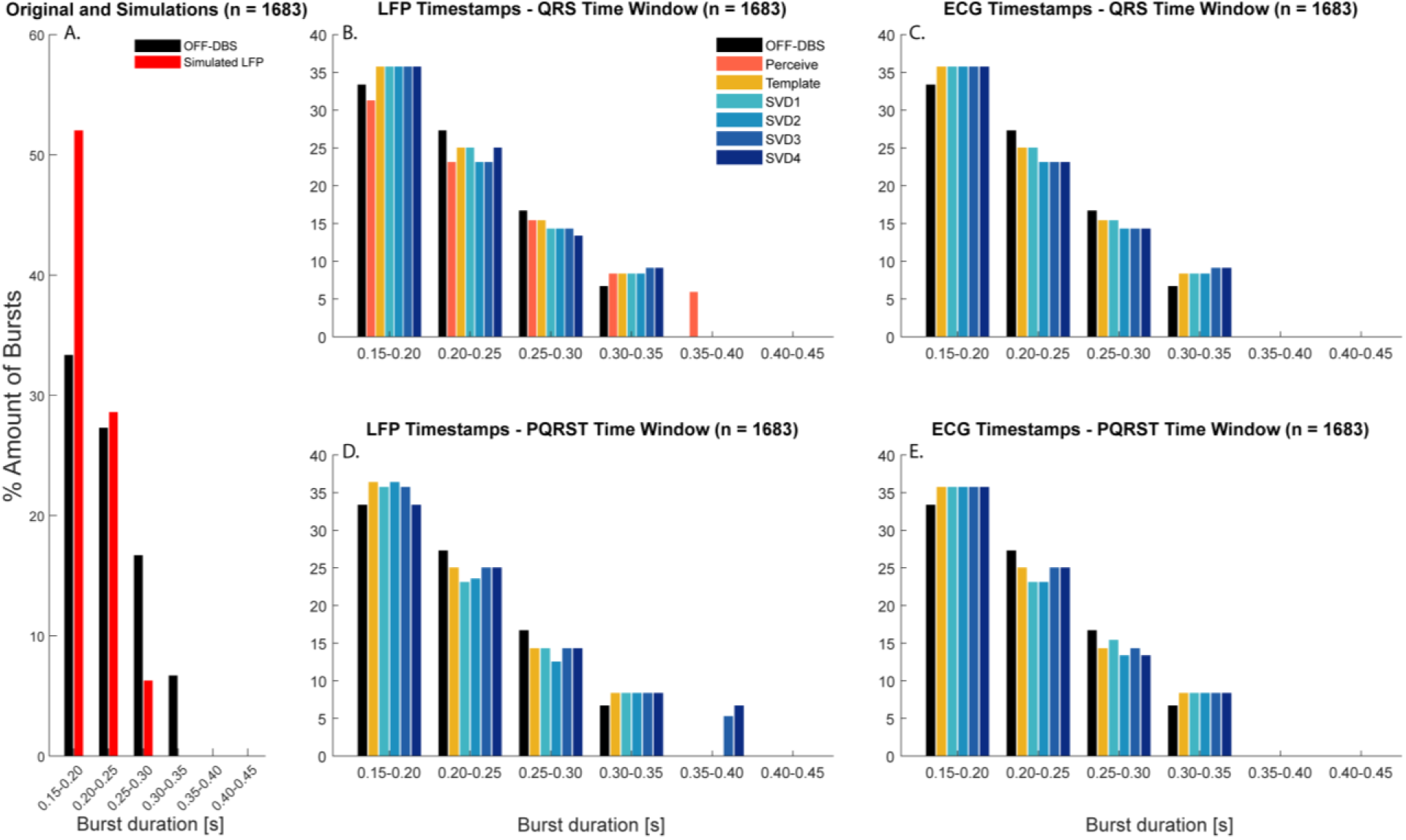
Distribution of median beta bursts durations. Beta burst duration distributions of the original OFF-DBS LFP signals and ECG artifact simulation signals, prior and after applying the ECG suppression methods. The distribution of burst durations was calculated as a percentage of total number of bursts of each simulation and averaged across 1683 simulations. Bursts were defined with a minimum duration of 150 ms and assigned to 50 ms bins.

**Table 3.**
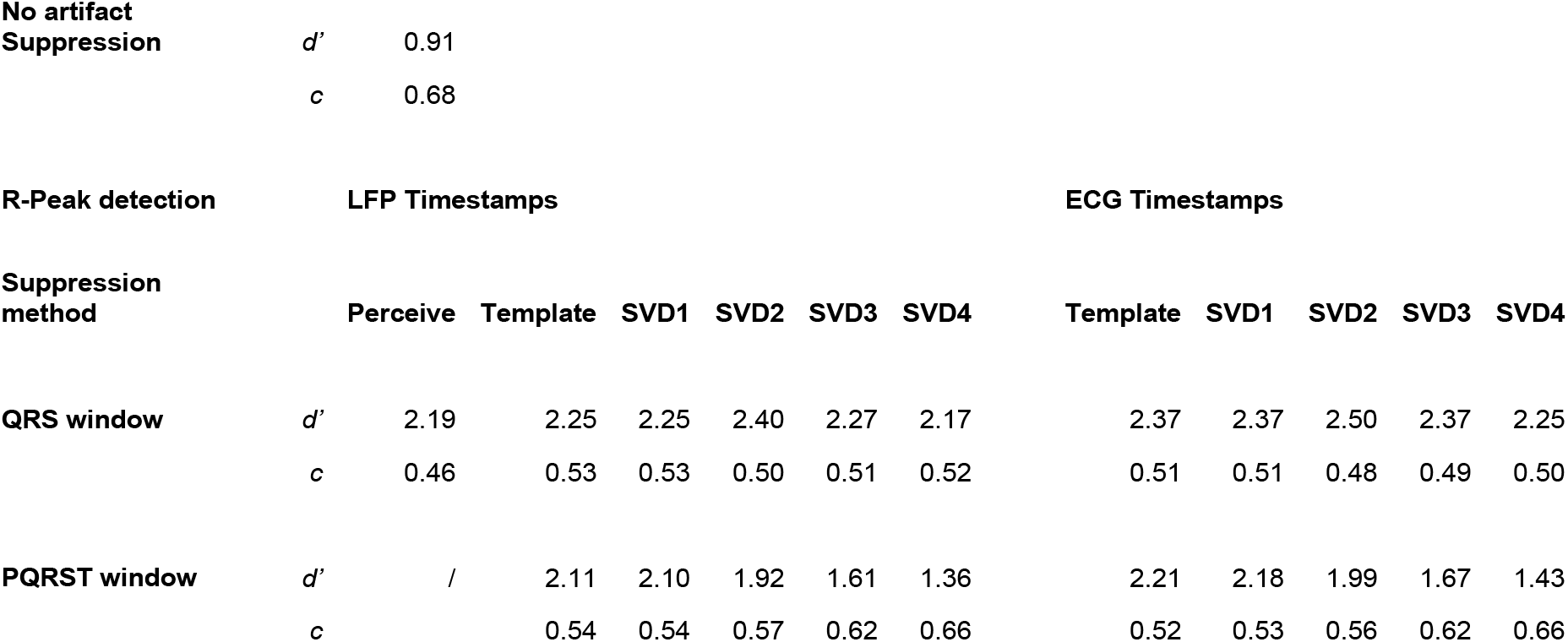
Accuracy of beta burst detection before and after applying the ECG suppression methods. The sensitivity index d’ and bias criterion c of the simulated LFP signals that contained a clear beta peak (n = 1683). For these results, all levels of ECG contamination were averaged. For better performance, sensitivity index d’ should go towards a value of 3 and bias criterion c towards 0.

The sensitivity index *d’* was calculated to express the ability of the ECG suppression methods to recover beta bursts. Table 3 suggest that the beta bursts are most accurately recovered after applying the SVD2QRS (ECG) method (see Table S4 in Supplementary Materials for the results divided into level of ECG contamination). The results suggest a better burst recovery when using the shorter QRS time window compared to the longer PQRST time window (*χ2*(1) = 1166.251, *p* < .001). In addition, using the timestamps of the externally recorded ECG signal to suppress ECG artifacts led to a slight improvement of beta burst sensitivity as compared to only using the LFP signal (*χ2*(1) = 62.760, *p* < .001). Dividing the results into level of ECG contamination showed that this slight improvement was mainly present at the lower levels of ECG contamination (p < 0.05 for 0-40; 60%). A significant higher *d’* for LFP timestamps was found at 80% ECG contamination (*χ2*(1) = 6.281, *p* = .012).

## Discussion

In this study, more than half of the ON-DBS 0 mA LFP signals recorded with the Medtronic PerceptTM PC system were contaminated with ECG artifacts. Three artifact-suppression methods were compared: a QRS interpolation as implemented in the Perceive toolbox, an optimized template subtraction method, and a singular value decomposition method. These methods were evaluated both in time- as well as in frequency domain. All evaluated ECG suppression methods were able to drastically reduce the percent difference of beta band power and at least partly recover the beta peak and beta burst dynamics. The SVD method seems most suited for artifact cleaning as long as its parameter settings are adequately chosen.

### Comparison of methods

Based on the data set used in this study, the SVD method was most suited for recovering beta band power. A benefit of applying the SVD method at the stage of post-processing is the possibility to tweak the settings around the R-peaks or select a different number of components. Although using a longer time window to create the ECG artifact epochs led to better reduction of beta band (and potentially low frequency) power difference, it increased the risk of flattening the beta peak and the loss of beta burst dynamics. Increasing the number of components might better capture any time-varying fluctuations in the PQRST waveform shape. The poorer performance of the template-based methods is possibly related to their inflexibility to do so. On the other hand, applying the SVDPQRST method with three or four components led to spectral patterns that were less similar to the corresponding/original OFF-DBS LFP signals, suggesting that relevant signal features were removed. Considering the different levels of ECG contamination in the simulated LFPs, recordings with large ECG artifacts could potentially benefit from increasing the time window and number of components of the SVD method. The negative percent differences in case of minor ECG artifact, on the other hand, suggests it might be better to use shorter time window and fewer components in LFPs with minor ECG artifacts. The optimal number of components might be difficult to determine based on the percentage of explained energy alone due to the large variability across subjects (Supplementary Materials; Table S2). For our data set, the SVD2QRS method formed the preferred trade-off between reducing percent difference of beta power and preserving the beta peak and bursts to study beta oscillations in Parkinson’s disease. Nevertheless, as the SVD method is designed for post-processing research analysis and developments of neuromodulation move towards adaptive DBS systems, the possibility of the template or Perceive methods to be applied automatically and in (semi) real time should be further explored.

Results showed that using a longer time window to create the ECG artifact epochs led to better reduction of beta band power difference. Increasing the time window means complementary suppression of the P- and T-waves of the ECG artifact, which was found to also better recover the spectral content of low frequencies (Hammer et al., 2022). Although not systematically assessed in this study, Figures 4 and 6 confirm the finding of Hammer et al. (2022) that the Perceive method less accurately suppresses low frequency power resulting from ECG remnants beyond the QRS peaks. An explanation for the poor suppression of low frequency content is that the neighbouring data points used for mirrored padding might contain a low frequency component of the ECG artifact, which is then duplicated. The QRS interpolation by mirrored padding was also the reason for not assessing the Perceive method with an increased time window, as this feature would have removed too much of the LFP signals.

### Visual inspection versus automatic peak detection

Neither of the two ECG detection algorithms assessed in this study identified all LFP signals with ECG artifact correctly, compared with visual inspection. Therefore, as yet, visual inspection of the LFP signals on the presence of ECG artifacts cannot be replaced by automatic ECG detection. In the future, machine learning may be used to detect ECG before a cleaning algorithm is applied (Merk et al., 2022). Additionally, no consensus between researchers could be reached about when exactly the R-peaks occurred in the LFP signals without using the externally recorded ECG signal to compare. Moreover, the occurrence of an R-peak in the external ECG signal does not necessarily serve as ground truth of the occurrence of an R-peak of the ECG artifact in the LFP signal. Whereas the current study ensured ECG artifact suppression by predetermining the R-peaks of the ECG artifact in the LFP signal using an external ECG signal, another solution is suggested by Chen and colleagues (2021). They demonstrated the feasibility of recording an ECG reference signal using the monopolar montage (between the IPG and remaining contact point) of a sensing-enabled neurostimulator. That method even allowed to extract a clear R-peak through large stimulation artifacts. In theory, the PerceptTM PC could incorporate such an ECG reference signal as well, but currently this is not implemented. The overall results of this study show that the performance of the ECG suppression methods improve by using the predetermined R-peaks. However, according to the simulations, in LFP signals with minor ECG contamination, the beta peak will still show if present and the artifact-induced increase of beta band power is only minimal. The application of ECG suppression methods may worsen the beta bursts detection and may result in negative percent differences. Ultimately, the conservative nature of the ECG detection algorithms and the fact that some R-peaks remain undetected when no additional ECG signal is used avoids the unnecessary loss of LFP information.

### Recovering beta bursts

Besides confirming the effect of the ECG suppression methods on restoring the beta peak and beta band power of the empirical data, the simulations also allow to quantitively study the effect on beta bursts dynamics in the time domain. The fact that all evaluated methods suppress ECG artifacts at the significantly affected segments of the LFP signal, instead of removing all these segments, is beneficial for the continuity of the recording. Cutting out the artifacts and attaching the rest of the recording or replacing the segments containing artifacts with zero would have made burst analysis impossible. Results show that ECG contamination led to a shift towards a higher number of bursts when contaminated by ECG. However, at the same time, the duration of the detected bursts tended to reduce. Results showed that applying the ECG suppression methods allowed to counter these shifts.

### Methodological considerations

ECG artifacts in the ON-DBS 0 mA LFP signals caused the difference in beta band power to greatly increase. However, care has to be taken when interpreting these results as our artifact-free recordings showed that solely turning on the Streaming mode at 0 mA already led to a difference in beta band power as compared to the OFF-DBS LFP signals. In addition, previous research found a variable scaling difference between ON-DBS 0 mA and OFF-DBS recordings for the same patient (Hammer et al., 2022). Apart from the contamination with ECG artifacts, the spectral difference between the ON-DBS 0 mA and OFF-DBS data can also be attributed to differences in patient state (even though the recordings were performed directly after another). Despite the limited recording montages available compared to the OFF-DBS mode, to avoid confounded differences in PSDs as a result of scaling differences, it is recommended to record ON-DBS 0 mA LFPs when collecting baseline recordings off stimulation.

### Cause of ECG artifacts

The ECG artifact can be attributed to an inadequate common mode rejection ratio (CMRR) of the sensing input chain of the DBS system. As LFP signals are measured as a differential signal, the ECG spikes are regarded as common mode signals which can be rejected by differentiating (Neumann et al., 2021). However, it is challenging to achieve a high CMRR in an implantable system. The size of the implant is greatly determined by the battery encapsulated in the case of the implanted pulse generator (IPG). As the size of the IPG is limited, so is its power consumption. To reduce power consumption, the PerceptTM PC uses passive recharge after the stimulation pulse for the redistribution of charge at the tissue-electrode interface. Passive instead of active recharge increases the duration the electrodes are connected to the IPG, while it is also the ground for the LFP recordings. This makes the recordings more susceptible for ECG contamination (Stanslaski et al., 2018). Additionally, slight leakage of fluid into the IPG can lead to an input impedance mismatch. Such mismatch breaks the symmetry of the differentiated signal, which alters the CMRR (Quinn et al., 2015).

### Recommendations for avoidance of ECG artifacts

Despite the ability of all ECG suppression methods to recover the beta peak and beta bursts, artifacts reduce the signal to noise ratio and hence the reliability of the LFPs for physiomarker and adaptive DBS research. Therefore, ECG artifacts should be avoided as much as possible. Suggested approaches to mitigate ECG contamination are improving the electrical properties of the electrode leads and extensions or developing new coatings to lower the impedance of the tissue-electrode interface. The use of a rechargeable battery could also mitigate ECG artifacts. DBS systems with a rechargeable battery can afford a higher power consumption, which would allow for active recharge after the stimulation pulse to be used. An alternative approach to reduce ECG artifacts is strategic placement of the IPG. Neumann and colleagues (2021) found that the ECG artifact appears to be stronger and more frequent when the IPG is implanted on the left side of the chest, when compared to right or cranial implants. The current study confirms that finding, as the Percept was implanted on the right side of the body in all LFP recordings that were free from ECG artifacts (Supplementary Materials; Table S1). Although a right chest/abdomen implant location reduces ECG contamination in LFP signals as compared to left implant locations, cranial mount systems might be even more robust to both ECG and movement artifact contamination. Further technological developments of DBS systems are important for future physiomarker and adaptive DBS research that aims to enhance the efficacy of DBS therapy based on reliable LFPs in which artifacts are minimized.

## Conclusion

DBS-LFP recordings are prone to ECG artifacts, and often require extensive cleaning before enabling further analysis of patient- and disease-specific neural activity. All of the assessed ECG suppression methods in this study were able to recover the beta peak and beta bursts. Based on the current data set, the SVD method was most suited for artifact cleaning as long as its parameter settings are adequately chosen. We recommend to adjust the epoch length around the R-peaks and the number of components based on the feature of the LFP signal that will be analysed.

## Data availability

The datasets generated during and analyzed during the current study are available from the corresponding author on reasonable request.

## Code availability

The code used to analyze data during the current study are available from the corresponding author on reasonable request.

## Acknowledgements

None.

---

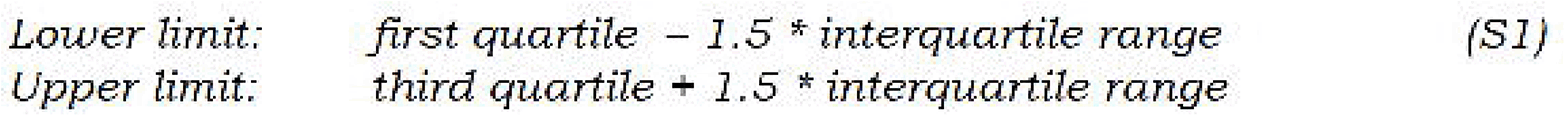

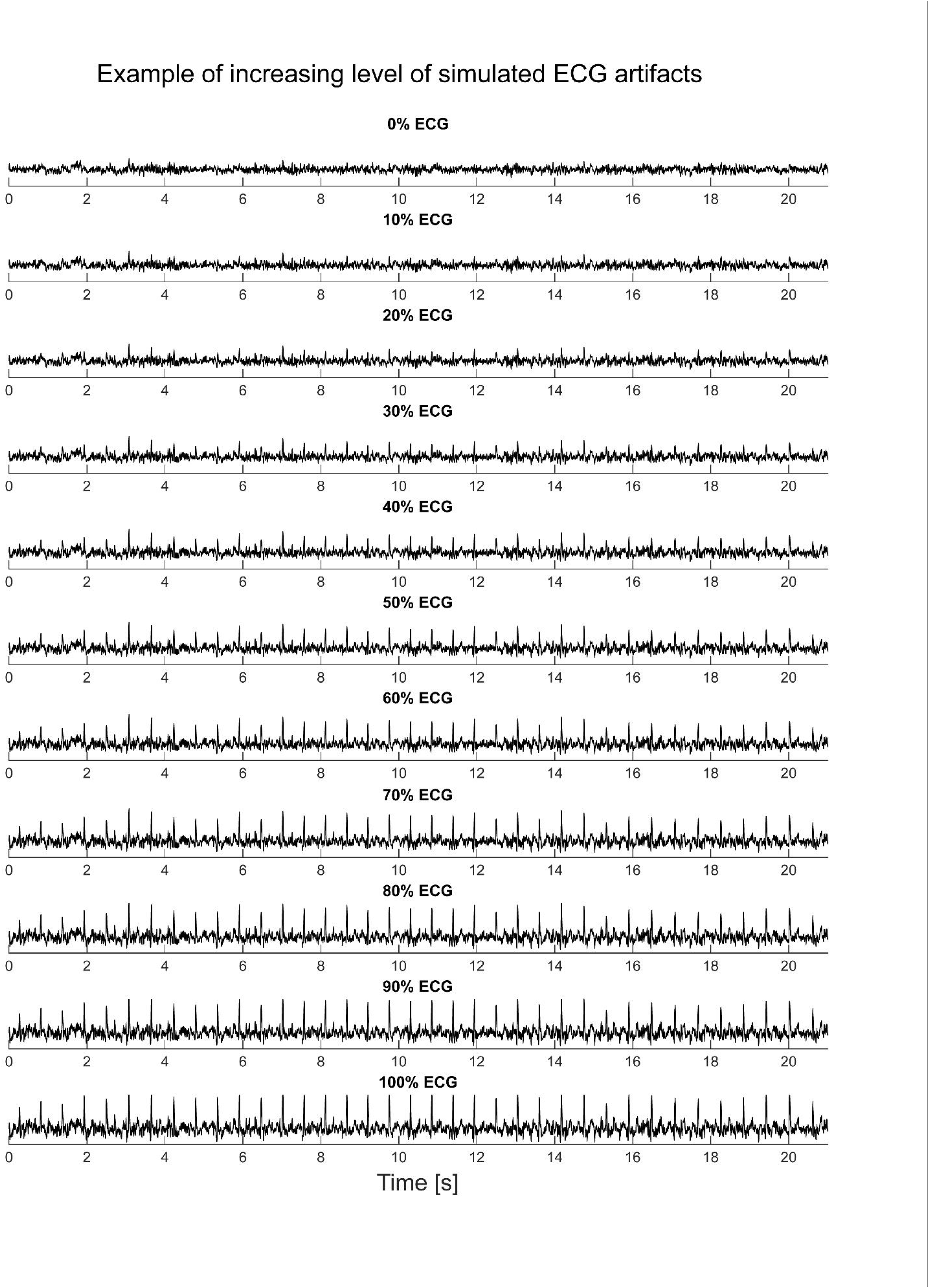

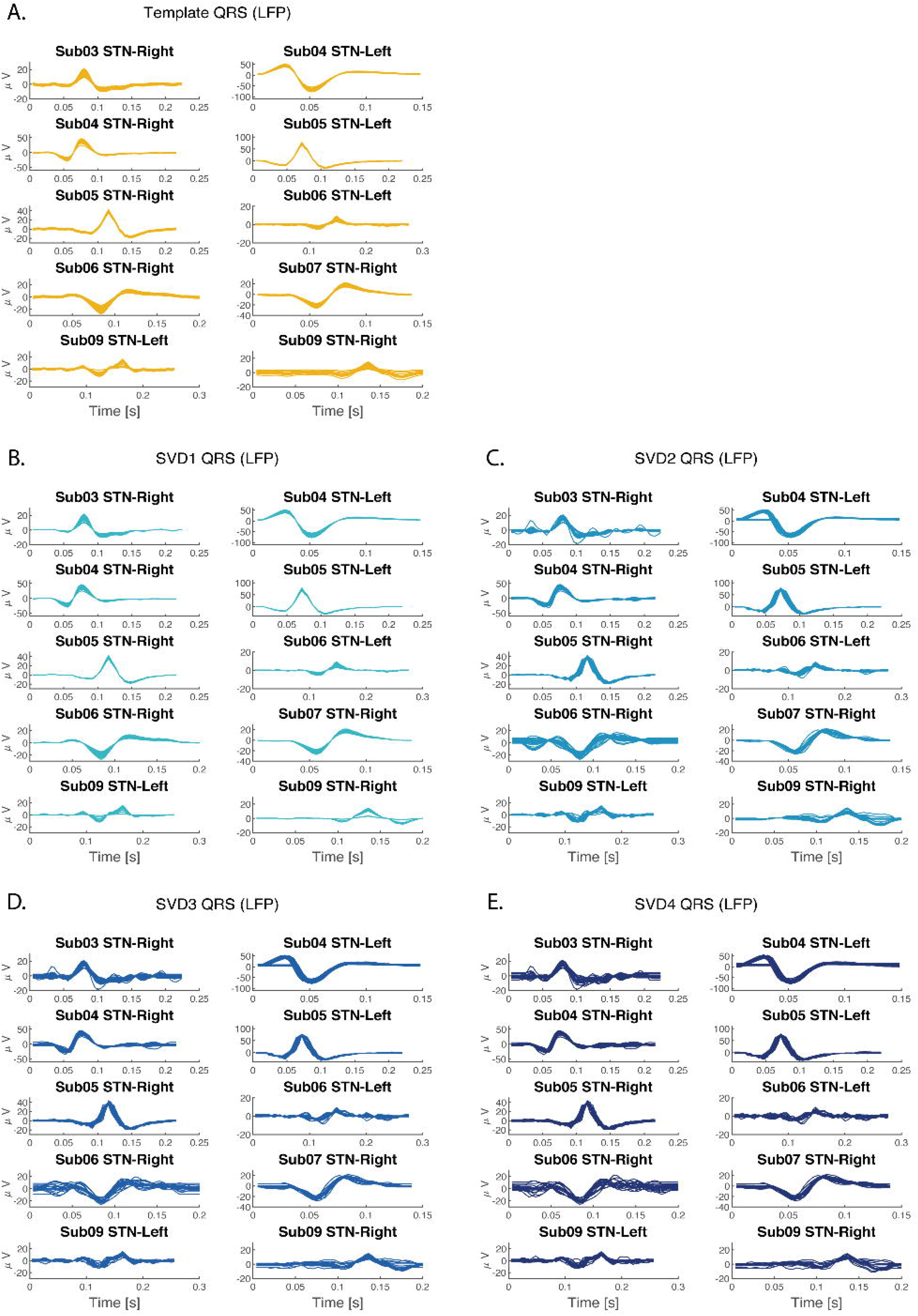

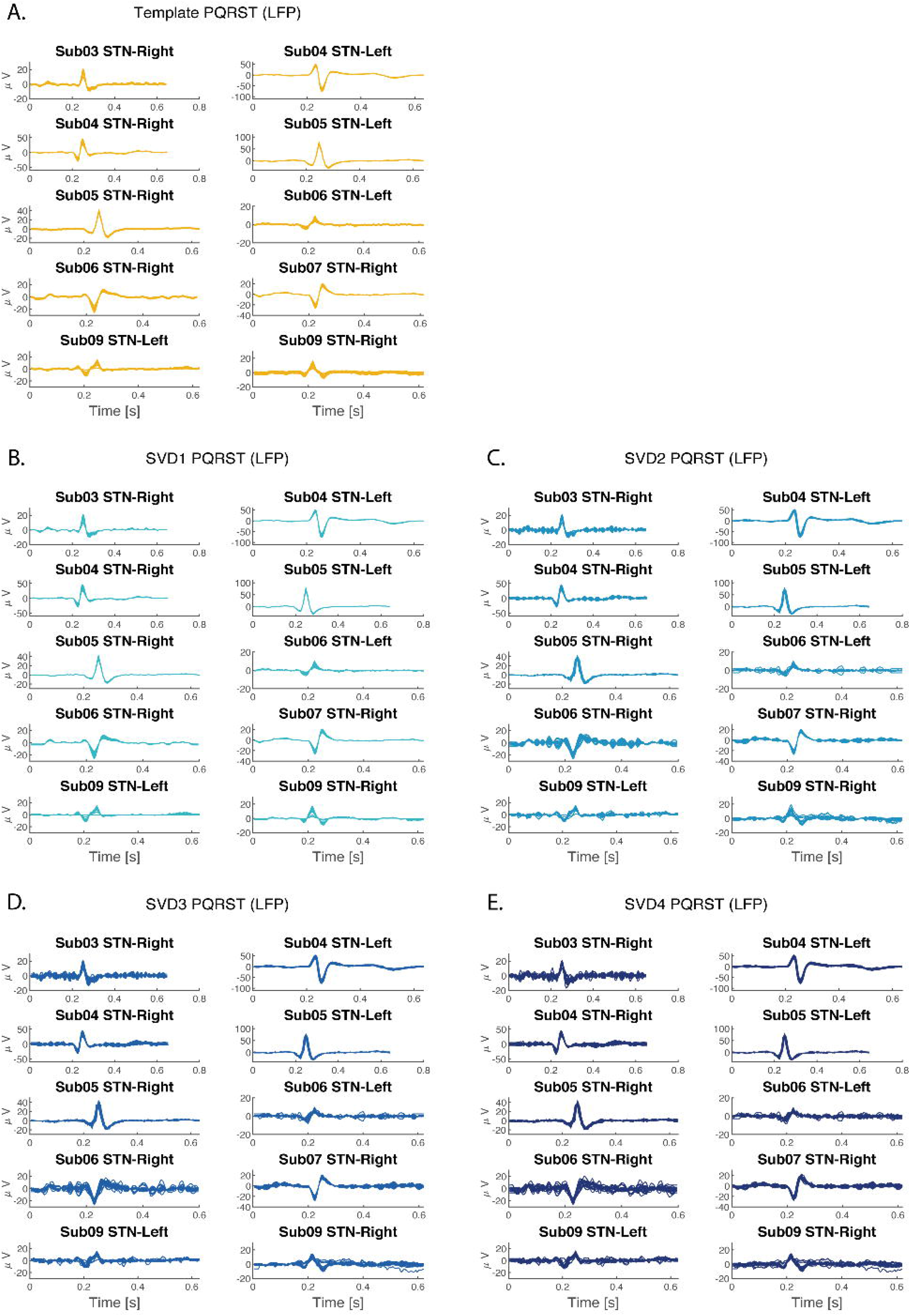

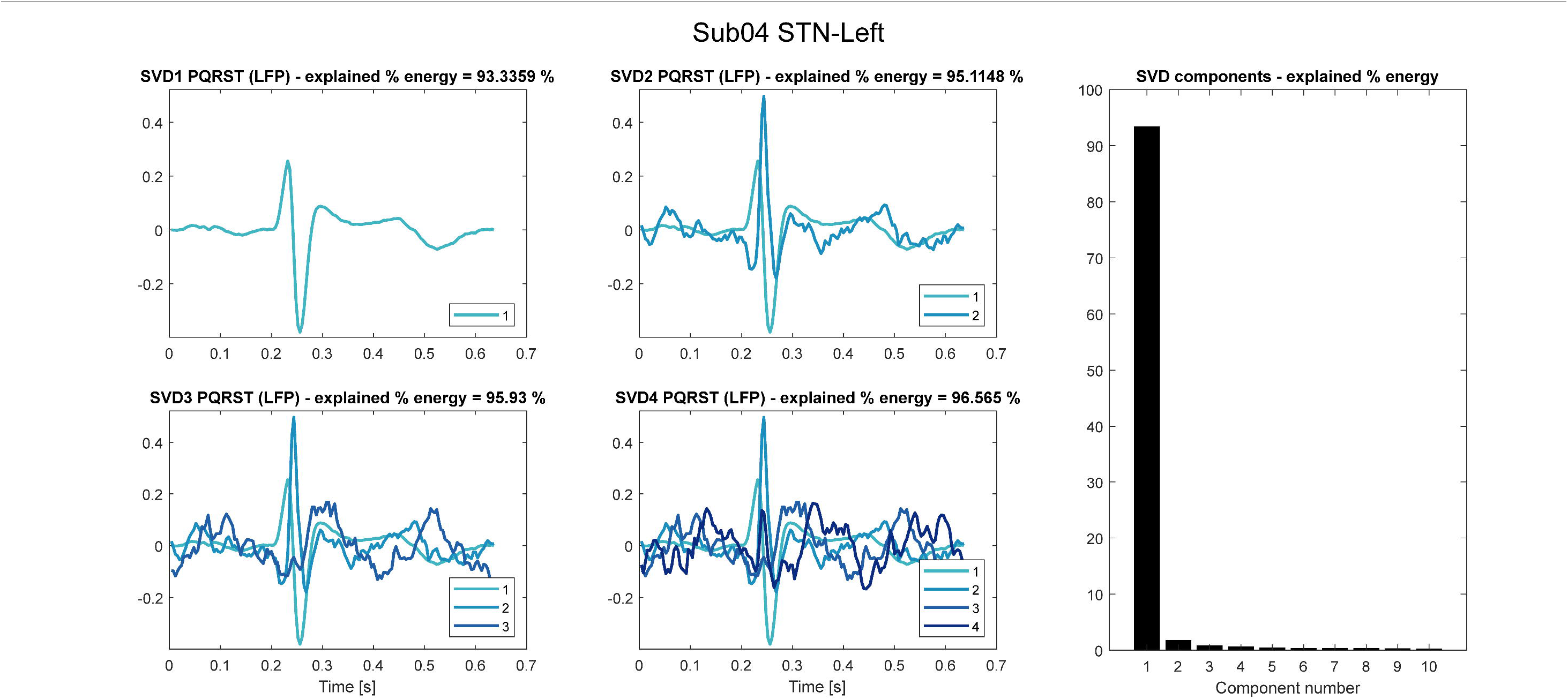

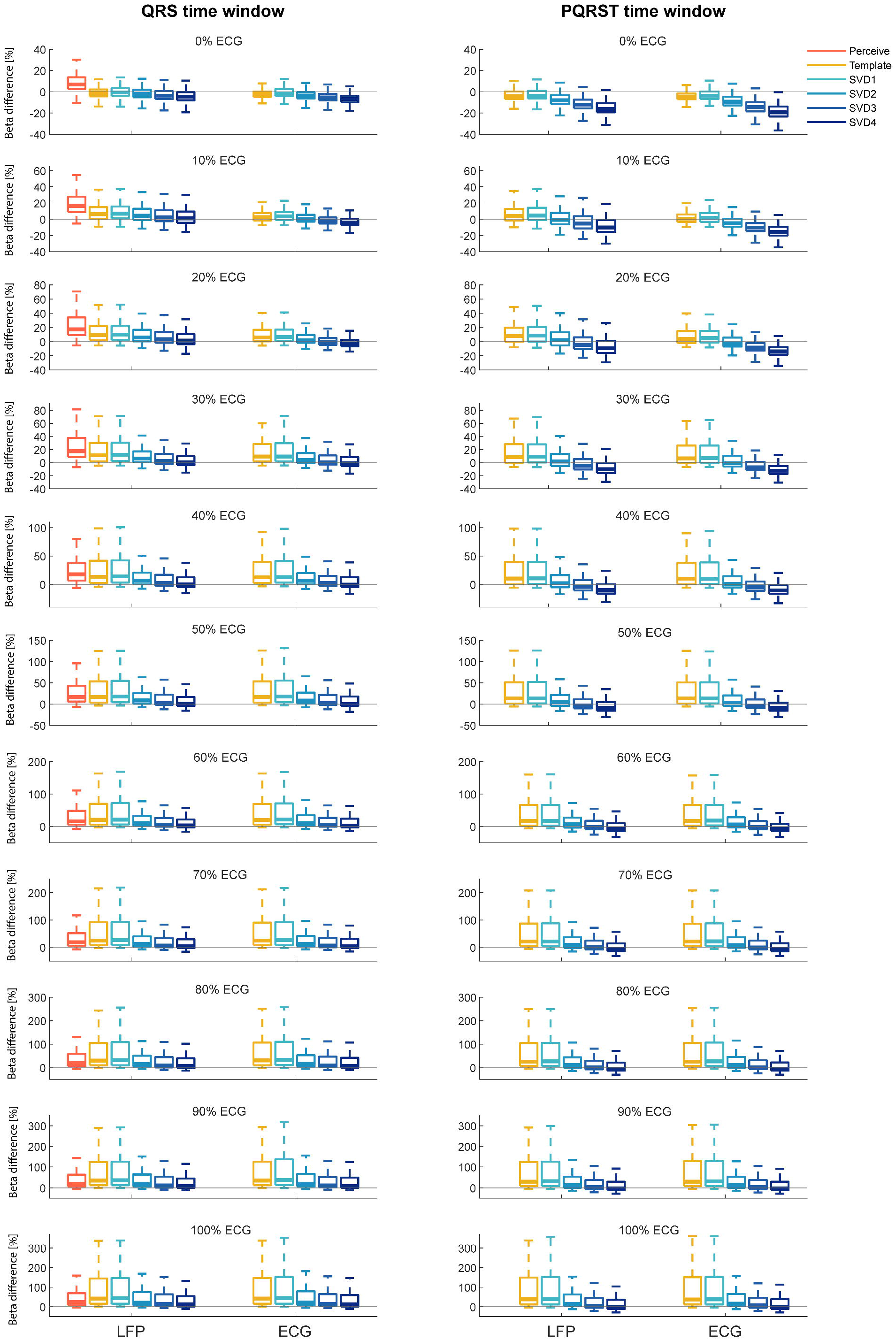

