## Supplementary Materials for "A comparison of methods to suppress electrocardiographic artifacts in local field potential recordings"

Table S1. Patient demographics.

| Subject | Age/Sex | IPG site | Lead Target | LFP Signal | ECG Artifact Y/N | Stimulation settings | Sensing Contacts |
|---|---|---|---|---|---|---|---|
| Sub01 | 59/F | Chest Right | STN-Left | LFP01 | N | C+9-; 0 mA; 130 Hz; 60 µs | 8 and 10 |
| | | | STN-Right | LFP02 | N | C+1-; 0 mA; 130 Hz; 50 µs | 0 and 2 |
| Sub02 | 54/M | Chest Right | STN-Left | LFP03 | N | C+9-; 0 mA; 130 Hz; 60 µs | 8 and 10 |
| | | | STN-Right | LFP04 | N | C+1-; 0 mA; 130 Hz; 60 µs | 0 and 2 |
| Sub03 | 52/F | Chest Right | STN-Left | LFP05 | N | C+9-; 0 mA; 130 Hz; 60 µs | 8 and 10 |
| | | | STN-Right | LFP06 | Y | C+2-; 0 mA; 130 Hz; 60 µs | 1 and 3 |
| Sub04 | 73/M | Chest Left | STN-Left | LFP07 | Y | C+9-; 0 mA; 180 Hz; 90 µs | 8 and 10 |
| | | | STN-Right | LFP08 | Y | C+2-; 0 mA; 180 Hz; 140 µs | 1 and 3 |
| Sub05 | 56/F | Chest Left | STN-Left | LFP09 | Y | C+10-; 0 mA; 130 Hz; 60 µs | 9 and 11 |
| | | | STN-Right | LFP10 | Y | C+2-; 0 mA; 130 Hz; 60 µs | 1 and 3 |
| Sub06 | 71/M | Chest Right | STN-Left | LFP11 | Y | C+9-; 0 mA; 125 Hz; 60 µs | 8 and 10 |
| | | | STN-Right | LFP12 | Y | C+1-; 0 mA; 125 Hz; 60 µs | 0 and 2 |
| Sub07 | 62/F | Abdomen Right | STN-Left | LFP13 | N | C+9-; 0 mA; 130 Hz; 60 µs | 8 and 10 |
| | | | STN-Right | LFP14 | Y | C+2-; 0 mA; 130 Hz; 60 µs | 1 and 3 |
| Sub08 | 52/F | Chest Right | STN-Left | LFP15 | N | C+9-; 0 mA; 130 Hz; 60 µs | 8 and 10 |
| | | | STN-Right | LFP16 | N | C+1-; 0 mA; 130 Hz; 60 µs | 0 and 2 |
| Sub09 | 63/M | Chest Left | STN-Left | LFP17 | Y | C+10-; 0 mA; 130 Hz; 60 µs | 9 and 11 |
| | | | STN-Right | LFP18 | Y | C+1-; 0 mA; 130 Hz; 60 µs | 0 and 2 |

Figure S1. An example of an OFF-DBS LFP signal added with eleven levels of simulated ECG artifact.

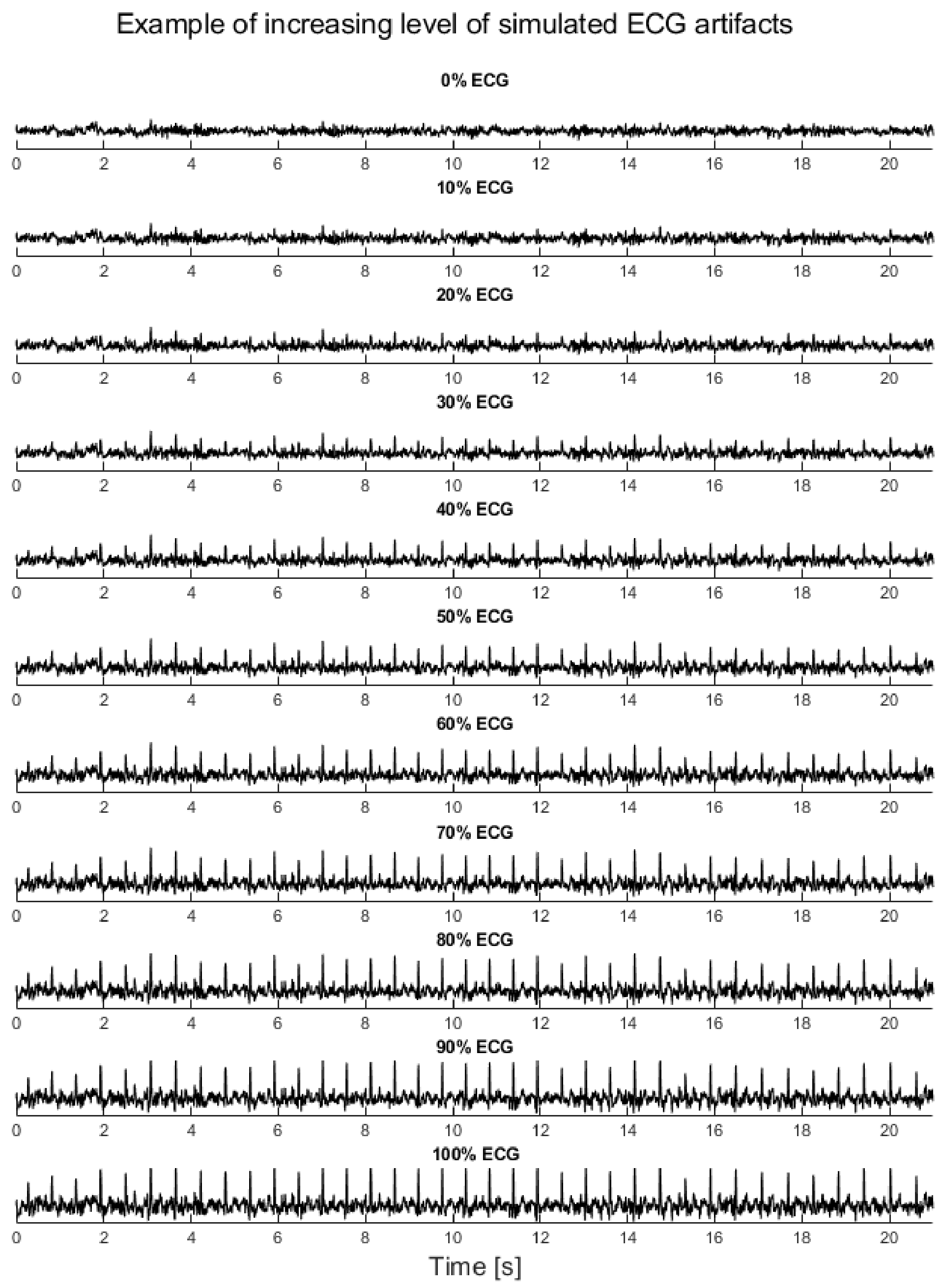

**Formula S1. Formula for determining adjacent values**. The lower and upper adjacent values are defined as the lowest and highest data points, respectively, that are still inside the region defined by the following limits:

$$\textit{Lower limit:} \quad \textit{first quartile} - 1.5 * \textit{interquartile range} \qquad (S1)$$
$$\textit{Upper limit:} \quad \textit{third quartile} + 1.5 * \textit{interquartile range}$$

**Figure S2. Artifact reconstructions for the Template (LFP) and SVD1-4 (LFP) methods with QRS time window.** Panel A shows the final QRS templates of the ten recorded LFP signals with ECG artifact. Per QRS epoch in the LFP signal, the final QRS template was scaled and the offset between the original R-peak and the template was corrected. Subsequently, the optimized template was subtracted from the original epoch to obtain a cleaned LFP signal. Panels B-E show the artifact reconstructions of the ten recorded LFP signals with ECG artifact for the SVD1-4 (LFP) methods. Each individual line in the plots indicate the component(s) detected for a single QRS epoch, which was subtracted from that specific epoch to obtain a cleaned LFP signal.

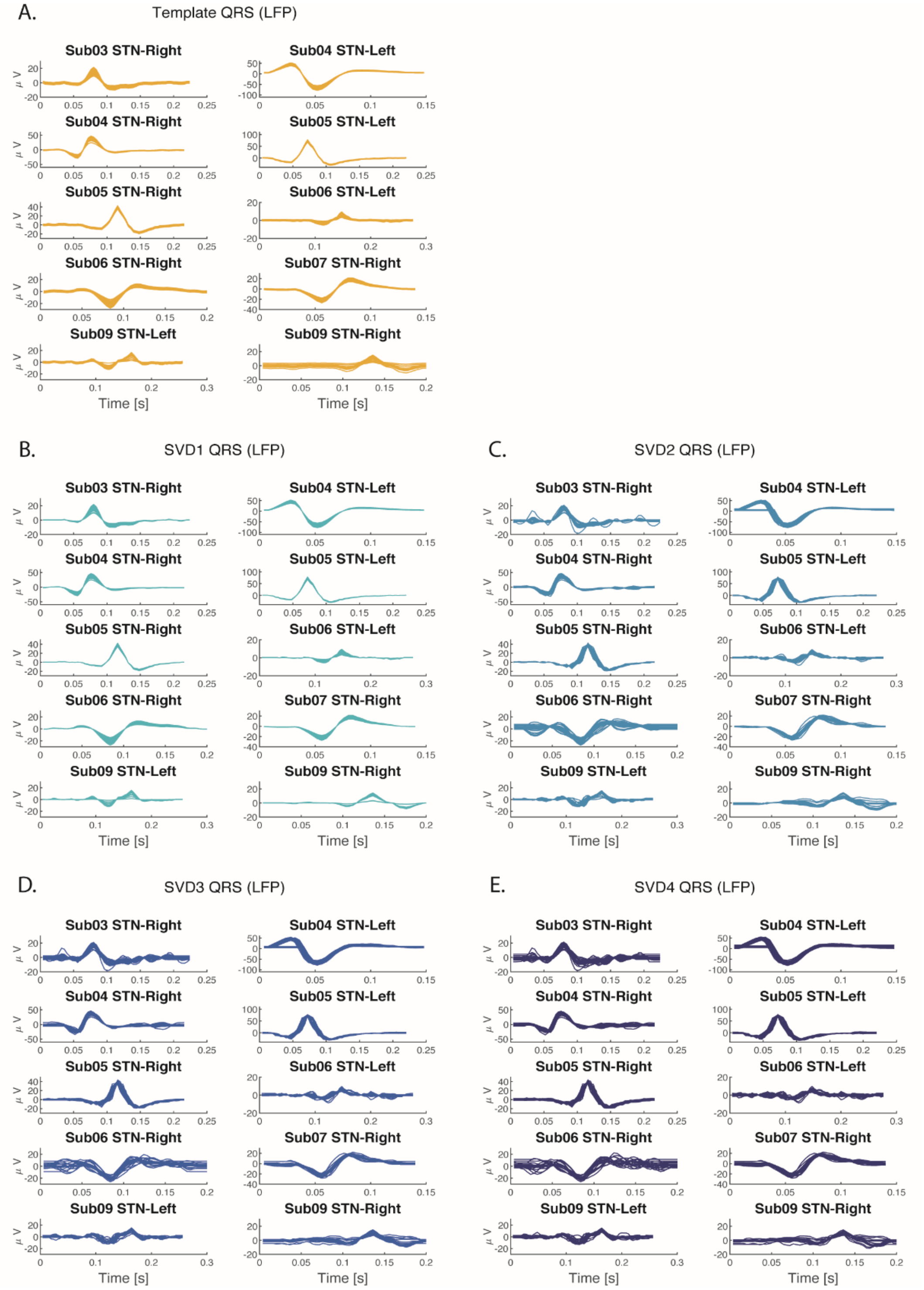

**Figure S3. Artifact reconstructions for the Template (LFP) and SVD1-4 (LFP) methods with PQRST time window.** Panel A shows the final PQRST templates of the ten recorded LFP signals with ECG artifact. Per PQRST epoch in the LFP signal, this final PQRST template was scaled and the offset between the original R-peak and the template was corrected. Subsequently, the optimized template was subtracted from the original epoch to obtain a cleaned LFP signal. Panels B-E show the artifact reconstructions of the ten recorded LFP signals with ECG artifact for the SVD1-4 (LFP) methods. Each individual line in the plots indicate the component(s) detected for a single PQRST epoch, which was subtracted from that specific epoch to obtain a cleaned LFP signal.

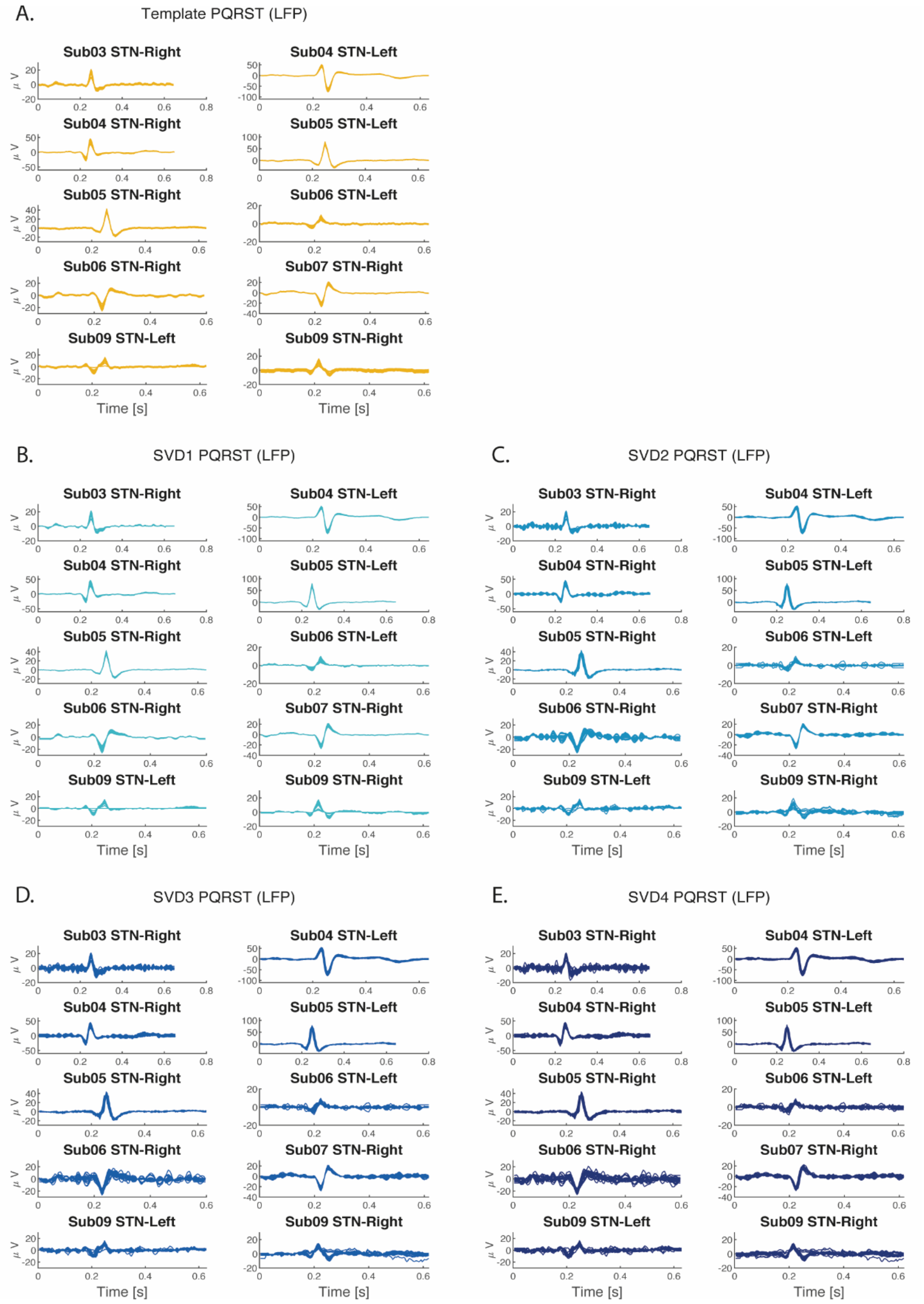

Table S2. The cumulative percentage energy of the components of the SVD methods.

| | QRS Window | | | | | | | |
|---|---|---|---|---|---|---|---|---|
| | LFP Timestamps | | | | ECG Timestamps | | | |
| [%] | SVD1 | SVD2 | SVD3 | SVD4 | SVD1 | SVD2 | SVD3 | SVD4 |
| **Sub03-R** | 68.248 | 75.091 | 80.239 | 84.306 | 64.058 | 71.684 | 76.854 | 81.099 |
| **Sub04-L** | 97.012 | 98.682 | 99.177 | 99.452 | 97.007 | 98.683 | 99.177 | 99.456 |
| **Sub04-R** | 89.232 | 91.813 | 93.785 | 95.494 | 89.100 | 91.724 | 93.696 | 95.439 |
| **Sub05-L** | 92.981 | 98.113 | 98.557 | 98.858 | 92.968 | 98.101 | 98.550 | 98.851 |
| **Sub05-R** | 92.130 | 96.271 | 97.189 | 97.770 | 92.137 | 96.278 | 97.187 | 97.772 |
| **Sub06-L** | 46.555 | 56.323 | 64.245 | 70.454 | 47.480 | 57.980 | 64.945 | 71.025 |
| **Sub06-R** | 64.300 | 77.479 | 82.790 | 87.451 | 57.422 | 69.847 | 76.737 | 81.115 |
| **Sub07-R** | 92.567 | 94.945 | 96.081 | 97.060 | 92.556 | 94.929 | 96.076 | 97.046 |
| **Sub09-L** | 63.382 | 71.857 | 78.618 | 82.907 | 61.242 | 69.805 | 75.580 | 79.573 |
| **Sub09-R** | 59.965 | 76.742 | 85.257 | 89.862 | 61.060 | 74.673 | 81.856 | 86.689 |

| | PQRST Window | | | | | | | |
|---|---|---|---|---|---|---|---|---|
| | LFP Timestamps | | | | ECG Timestamps | | | |
| [%] | SVD1 | SVD2 | SVD3 | SVD4 | SVD1 | SVD2 | SVD3 | SVD4 |
| **Sub03-R** | 44.348 | 51.003 | 57.129 | 62.249 | 43.000 | 49.660 | 55.932 | 60.646 |
| **Sub04-L** | 93.336 | 95.115 | 95.930 | 96.565 | 93.350 | 95.128 | 95.942 | 96.583 |
| **Sub04-R** | 73.351 | 77.600 | 81.115 | 83.763 | 73.328 | 77.576 | 81.126 | 83.783 |
| **Sub05-L** | 90.859 | 95.900 | 96.480 | 96.921 | 90.856 | 95.896 | 96.476 | 96.917 |
| **Sub05-R** | 88.222 | 92.344 | 93.447 | 94.276 | 88.217 | 92.337 | 93.440 | 94.286 |
| **Sub06-L** | 30.264 | 39.722 | 48.064 | 54.186 | 26.260 | 34.075 | 40.903 | 46.562 |
| **Sub06-R** | 39.842 | 50.765 | 59.714 | 65.550 | 35.873 | 45.002 | 53.218 | 59.322 |
| **Sub07-R** | 74.209 | 78.124 | 81.142 | 83.626 | 74.248 | 78.162 | 81.179 | 83.647 |
| **Sub09-L** | 44.570 | 52.622 | 59.283 | 65.271 | 48.099 | 54.898 | 61.508 | 66.835 |
| **Sub09-R** | 36.316 | 51.744 | 64.307 | 72.116 | 38.715 | 53.242 | 62.950 | 69.734 |

**Figure S4. The SVD components and corresponding explained percent energy of the left STN-LFP signal of Sub04.** In this example, the components were created using the LFP timestamps and the PQRST time window.

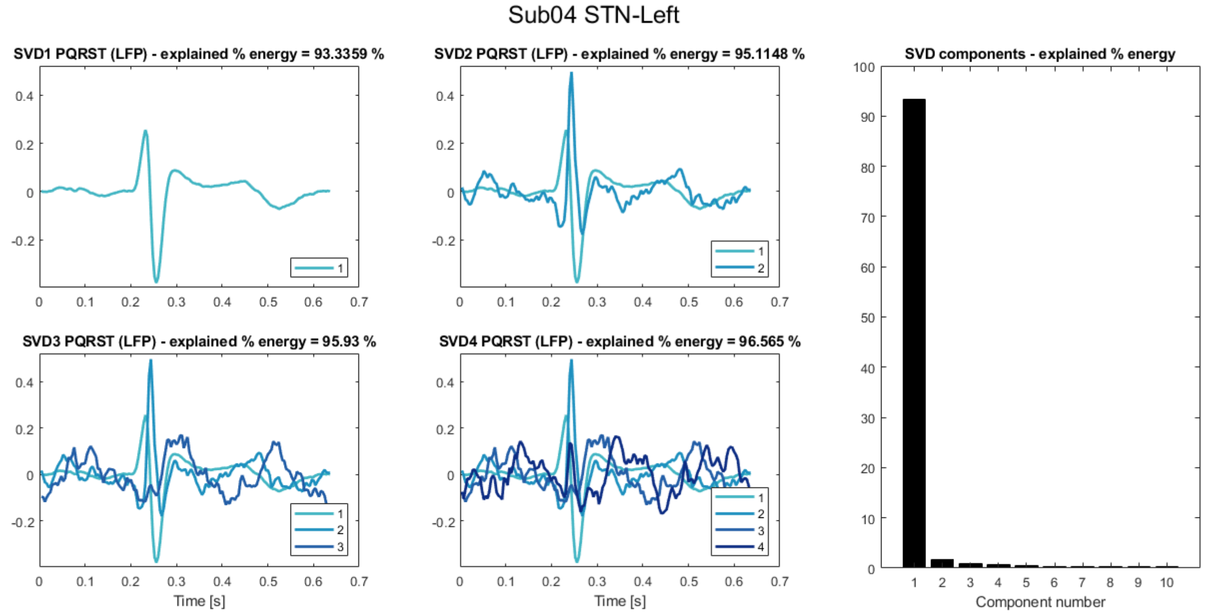

**Figure S5. Percentage difference of beta band power between the simulated LFP signals and corresponding OFF-DBS LFP signal, before and after applying the ECG suppression methods.** The simulated LFPs are divided into the different levels of ECG contamination (*n* = 486).

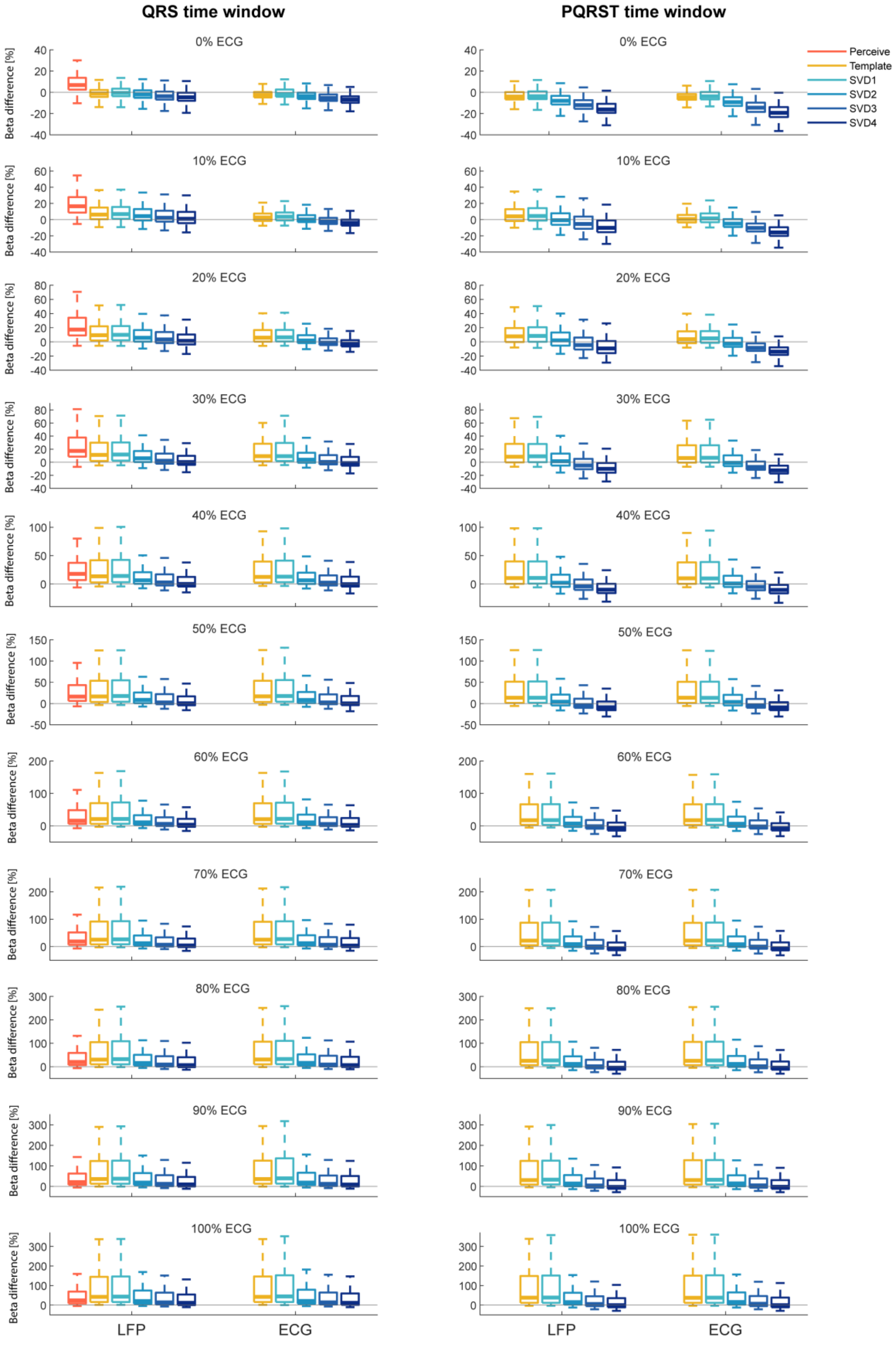

**Table S3. Duration and length of beta bursts**. The number of beta bursts and their duration (in ms) found in the original OFF-DBS LFP signals and the simulated LFPs with ECG artifact (*n* = 1683), prior and after applying the ECG suppression methods. For these results, all levels of ECG contamination were averaged.

| *N = 1683* | Number of Bursts (mean ± sd) | Duration of Bursts (mean ± sd ms) |
|---|---|---|
| Original LFPs | 12.12 ± 1.97 | 268 2 ± 27.8 |
| Simulations | 18.72 ± 4.88 | 215.8 ± 32.4 |
| Perceive | 12.81 ± 1.92 | 272.7 ± 32.5 |
| Template QRS (LFP) | 12.81 ± 2.02 | 256.4 ± 28.5 |
| Template QRS (ECG) | 12.83 ± 2.02 | 257.3 ± 28.4 |
| SVD1 QRS (LFP) | 12.82 ± 2.04 | 256.6 ± 28.2 |
| SVD1 QRS (ECG) | 12.86 ± 2.14 | 257.4 ± 28.6 |
| SVD2 QRS (LFP) | 12.59 ± 1.91 | 259.3 ± 28.8 |
| SVD2 QRS (ECG) | 12.67 ± 1.97 | 258.9 ± 28.8 |
| SVD3 QRS (LFP) | 12.71 ± 1.89 | 257.4 ± 29.2 |
| SVD3 QRS (ECG) | 12.79 ± 1.88 | 257.7 ± 28.5 |
| SVD4 QRS (LFP) | 12.77 ± 1.86 | 257.3 ± 28.2 |
| SVD4 QRS (ECG) | 12.92 ± 1.84 | 256.6 ± 26.9 |
| Template PQRST (LFP) | 12.84 ± 2.06 | 256.6 ± 31.5 |
| Template PQRST (ECG) | 12.86 ± 2.04 | 257.5 ± 31.8 |
| SVD1 PQRST (LFP) | 12.85 ± 2.02 | 257.1 ± 30.7 |
| SVD1 PQRST (ECG) | 12.82 ± 1.99 | 258.1 ± 31.2 |
| SVD2 PQRST (LFP) | 12.46 ± 1.87 | 260.5 ± 33.3 |
| SVD2 PQRST (ECG) | 12.51 ± 1.88 | 259.1 ± 32.7 |
| SVD3 PQRST (LFP) | 12.52 ± 1.92 | 259.8 ± 31.4 |
| SVD3 PQRST (ECG) | 12.53 ± 1.96 | 257.7 ± 30.3 |
| SVD4 PQRST (LFP) | 12.66 ± 1.94 | 260.5 ± 32.0 |
| SVD4 PQRST (ECG) | 12.65 ± 1.98 | 256.4 ± 30.5 |

**Table S4. Accuracy of beta burst detection before and after applying the ECG suppression methods.** The sensitivity index *d′* and bias criterion *c* of the simulated LFP signals that contained a clear beta peak. For these results, the simulated LFPs are divided into the different levels of ECG contamination (*n* = 153). For better performance, sensitivity index *d′* should go towards a value of 3 and bias criterion *c* towards 0.

| R-Peak detection | | | | LFP Timestamps | | | | | | ECG Timestamps | | | | |
|---|---|---|---|---|---|---|---|---|---|---|---|---|---|---|
| Method (based on QRS window) | | Original | Sim | Perceive | Template | SVD1 | SVD2 | SVD3 | SVD4 | Template | SVD1 | SVD2 | SVD3 | SVD4 |
| *0%* | *d′* | Inf | 3.30 | 2.36 | 2.24 | 2.31 | 2.09 | 2.01 | 1.96 | 2.72 | 2.78 | 2.52 | 2.33 | 2.25 |
| | *c* | NaN | 0.80 | 0.55 | 0.61 | 0.62 | 0.60 | 0.60 | 0.59 | 0.58 | 0.58 | 0.57 | 0.56 | 0.57 |
| *10%* | *d′* | Inf | 2.18 | 2.09 | 2.38 | 2.39 | 2.26 | 2.16 | 2.10 | 2.82 | 2.83 | 2.65 | 2.45 | 2.35 |
| | *c* | NaN | 0.70 | 0.43 | 0.53 | 0.53 | 0.55 | 0.55 | 0.56 | 0.50 | 0.50 | 0.52 | 0.54 | 0.55 |
| *20%* | *d′* | Inf | 1.45 | 2.16 | 2.57 | 2.57 | 2.49 | 2.37 | 2.24 | 2.80 | 2.81 | 2.72 | 2.53 | 2.35 |
| | *c* | NaN | 0.65 | 0.42 | 0.49 | 0.50 | 0.50 | 0.49 | 0.50 | 0.45 | 0.46 | 0.45 | 0.45 | 0.46 |
| *30%* | *d′* | Inf | 0.98 | 2.22 | 2.59 | 2.58 | 2.61 | 2.42 | 2.31 | 2.70 | 2.69 | 2.72 | 2.51 | 2.38 |
| | *c* | NaN | 0.70 | 0.51 | 0.57 | 0.57 | 0.53 | 0.56 | 0.54 | 0.54 | 0.54 | 0.48 | 0.51 | 0.51 |
| *40%* | *d′* | Inf | 0.66 | 2.33 | 2.50 | 2.50 | 2.60 | 2.43 | 2.30 | 2.55 | 2.53 | 2.64 | 2.49 | 2.34 |
| | *c* | NaN | 0.72 | 0.33 | 0.63 | 0.63 | 0.39 | 0.40 | 0.40 | 0.62 | 0.62 | 0.37 | 0.38 | 0.37 |
| *50%* | *d′* | Inf | 0.46 | 2.29 | 2.39 | 2.38 | 2.56 | 2.42 | 2.29 | 2.42 | 2.41 | 2.60 | 2.46 | 2.28 |
| | *c* | NaN | 0.61 | 0.42 | 0.43 | 0.43 | 0.42 | 0.44 | 0.47 | 0.42 | 0.42 | 0.44 | 0.44 | 0.45 |
| *60%* | *d′* | Inf | 0.33 | 2.24 | 2.24 | 2.22 | 2.49 | 2.35 | 2.23 | 2.26 | 2.24 | 2.49 | 2.38 | 2.27 |
| | *c* | NaN | 0.68 | 0.43 | 0.43 | 0.42 | 0.44 | 0.47 | 0.50 | 0.41 | 0.42 | 0.44 | 0.46 | 0.50 |
| *70%* | *d′* | Inf | 0.24 | 2.18 | 2.14 | 2.12 | 2.44 | 2.31 | 2.20 | 2.14 | 2.12 | 2.46 | 2.35 | 2.23 |
| | *c* | NaN | 0.66 | 0.47 | 0.55 | 0.54 | 0.50 | 0.53 | 0.55 | 0.51 | 0.51 | 0.50 | 0.50 | 0.51 |
| *80%* | *d′* | Inf | 0.18 | 2.11 | 2.03 | 2.03 | 2.33 | 2.20 | 2.11 | 2.03 | 2.00 | 2.30 | 2.20 | 2.11 |
| | *c* | NaN | 0.61 | 0.55 | 0.62 | 0.61 | 0.59 | 0.61 | 0.61 | 0.61 | 0.61 | 0.57 | 0.58 | 0.59 |
| *90%* | *d′* | Inf | 0.14 | 2.06 | 1.88 | 1.86 | 2.26 | 2.15 | 2.07 | 1.89 | 1.86 | 2.21 | 2.19 | 2.07 |
| | *c* | NaN | 0.68 | 0.47 | 0.52 | 0.53 | 0.53 | 0.53 | 0.55 | 0.50 | 0.51 | 0.51 | 0.50 | 0.51 |
| *100%* | *d′* | Inf | 0.12 | 2.03 | 1.80 | 1.79 | 2.23 | 2.18 | 2.08 | 1.80 | 1.77 | 2.23 | 2.19 | 2.07 |
| | *c* | NaN | 0.61 | 0.43 | 0.48 | 0.49 | 0.45 | 0.46 | 0.47 | 0.45 | 0.47 | 0.44 | 0.44 | 0.44 |

| | | | | | | | | | | | | | |
|---|---|---|---|---|---|---|---|---|---|---|---|---|---|
| Average | *d′* | / | 0.91 | 2.19 | 2.25 | 2.25 | 2.40 | 2.27 | 2.17 | 2.37 | 2.37 | 2.50 | 2.37 | 2.25 |
| | *c* | | 0.68 | 0.46 | 0.53 | 0.53 | 0.50 | 0.51 | 0.52 | 0.51 | 0.51 | 0.48 | 0.49 | 0.50 |

| Method (based on PQRST window) | | Original | Sim | Perceive | Template | SVD1 | SVD2 | SVD3 | SVD4 | Template | SVD1 | SVD2 | SVD3 | SVD4 |
|---|---|---|---|---|---|---|---|---|---|---|---|---|---|---|
| *0%* | *d′* | Inf | / | / | 2.02 | 2.06 | 1.54 | 1.23 | 1.09 | 2.42 | 2.33 | 1.78 | 1.48 | 1.28 |
| | *c* | NaN | | | 0.63 | 0.63 | 0.67 | 0.72 | 0.77 | 0.61 | 0.62 | 0.67 | 0.72 | 0.77 |
| *10%* | *d′* | Inf | / | / | 2.14 | 2.11 | 1.70 | 1.40 | 1.17 | 2.49 | 2.37 | 1.92 | 1.59 | 1.39 |
| | *c* | NaN | | | 0.54 | 0.55 | 0.63 | 0.69 | 0.73 | 0.52 | 0.54 | 0.63 | 0.70 | 0.73 |
| *20%* | *d′* | Inf | / | / | 2.31 | 2.26 | 1.87 | 1.58 | 1.31 | 2.52 | 2.45 | 2.04 | 1.72 | 1.47 |
| | *c* | NaN | | | 0.51 | 0.50 | 0.59 | 0.65 | 0.67 | 0.48 | 0.48 | 0.58 | 0.65 | 0.68 |
| *30%* | *d′* | Inf | / | / | 2.36 | 2.33 | 2.03 | 1.69 | 1.42 | 2.43 | 2.39 | 2.09 | 1.76 | 1.50 |
| | *c* | NaN | | | 0.59 | 0.58 | 0.60 | 0.63 | 0.67 | 0.57 | 0.57 | 0.59 | 0.63 | 0.67 |
| *40%* | *d′* | Inf | / | / | 2.30 | 2.26 | 2.06 | 1.72 | 1.43 | 2.33 | 2.31 | 2.07 | 1.75 | 1.45 |
| | *c* | NaN | | | 0.63 | 0.62 | 0.52 | 0.57 | 0.60 | 0.63 | 0.62 | 0.52 | 0.57 | 0.59 |
| *50%* | *d′* | Inf | / | / | 2.24 | 2.21 | 2.04 | 1.70 | 1.44 | 2.26 | 2.24 | 2.07 | 1.72 | 1.46 |
| | *c* | NaN | | | 0.42 | 0.42 | 0.51 | 0.56 | 0.62 | 0.41 | 0.43 | 0.51 | 0.58 | 0.62 |
| *60%* | *d′* | Inf | / | / | 2.15 | 2.13 | 2.01 | 1.70 | 1.42 | 2.17 | 2.15 | 2.03 | 1.70 | 1.43 |
| | *c* | NaN | | | 0.45 | 0.46 | 0.55 | 0.64 | 0.69 | 0.43 | 0.45 | 0.54 | 0.64 | 0.71 |
| *70%* | *d′* | Inf | / | / | 2.05 | 2.04 | 1.97 | 1.68 | 1.42 | 2.06 | 2.06 | 2.01 | 1.69 | 1.43 |
| | *c* | NaN | | | 0.55 | 0.55 | 0.56 | 0.63 | 0.67 | 0.51 | 0.53 | 0.56 | 0.63 | 0.69 |
| *80%* | *d′* | Inf | / | / | 1.97 | 1.97 | 1.98 | 1.67 | 1.43 | 1.98 | 1.96 | 2.00 | 1.67 | 1.43 |
| | *c* | NaN | | | 0.61 | 0.61 | 0.62 | 0.65 | 0.67 | 0.59 | 0.59 | 0.60 | 0.63 | 0.67 |
| *90%* | *d′* | Inf | / | / | 1.89 | 1.89 | 1.97 | 1.69 | 1.42 | 1.90 | 1.89 | 1.96 | 1.67 | 1.43 |
| | *c* | NaN | | | 0.54 | 0.53 | 0.54 | 0.59 | 0.62 | 0.51 | 0.51 | 0.51 | 0.56 | 0.60 |
| *100%* | *d′* | Inf | / | / | 1.79 | 1.79 | 1.93 | 1.67 | 1.42 | 1.79 | 1.79 | 1.93 | 1.66 | 1.44 |
| | *c* | NaN | | | 0.51 | 0.50 | 0.52 | 0.52 | 0.53 | 0.48 | 0.48 | 0.49 | 0.49 | 0.51 |
| Average | *d′* | / | / | / | 2.11 | 2.10 | 1.92 | 1.61 | 1.36 | 2.21 | 2.18 | 1.99 | 1.67 | 1.43 |
| | *c* | | | | 0.54 | 0.54 | 0.57 | 0.62 | 0.66 | 0.52 | 0.53 | 0.56 | 0.62 | 0.66 |
